# Whole-genome sequencing in Galicia reveals male-biased pre-Islamic North African ancestry, subtle population structure, and micro-geographic patterns of disease risk

**DOI:** 10.1101/2025.06.27.662083

**Authors:** Jacobo Pardo-Seco, Alba Camino-Mera, Xabier Bello, Alberto Gómez-Carballa, Lúa Castelo-Martínez, Conrado Martinez-Cadenas, Federico Martinón-Torres, Antonio Salas, the GALOMICS Working Group

## Abstract

Galicia, located at the westernmost edge of Europe, has been reported to exhibit distinctive genetic traits compared to other Iberian populations. We present the first whole-genome sequencing (WGS) study of a Galician population (GALOMICS [GAL]; *n* = 91, 17.2 million variants; https://galomica.genpob.eu/), assessing its genetic variability in comparison with WGS data from other Spanish regions and continental populations (*n* = 1,078). Contrary to recent claims of extreme genetic stratification, the population structure of Galicia aligns with broader Iberian patterns, with one dominant cluster homogeneously distributed and four minor, more localized clusters. Genome-wide analyses of the Spanish National DNA Bank dataset (NDNAB; *n* = 453) support these findings, identifying only three Galician clusters, again with one overwhelmingly predominant. Phylogenetic analyses challenge earlier interpretations that placed Galicians at the deepest Iberian genetic node; instead, Galician clusters form terminal branches, suggesting recent diversification. Analysis of runs of homozygosity indicates slightly higher inbreeding compared to other European populations, primarily driven by the ‘Porto do Son’ cluster, which raises the regional average. We identified a significant North African/Middle Eastern autosomal ancestry component (13.5%–16.5%), despite its distance from historically Arab-influenced regions. Genomic analyses point to an admixture event ca. 620–670 CE that introduced North African/Middle Eastern ancestry into a largely European gene pool. The signal, likely stemming from trans-Mediterranean contacts predating the 711 CE Islamic incursion and well before the *Reconquista*, shows a subtle South-to-North decline, suggesting a southern entry route. Y-chromosome (21.2%) and mitochondrial DNA (1.1%) analyses indicate a male-biased influx, pointing to a predominantly paternal contribution. This observation calls for a reevaluation of the commonly held assumption that Islamic rule alone accounts for North African ancestry in Iberia. Analysis of polygenic risk scores for common diseases (including breast and ovarian cancer, Alzheimer’s, schizophrenia, and type 2 diabetes) reveals distinct micro-geographical patterns of disease risk in the region, stratified by genetic clusters. These insights highlight the importance of further research into implications for public health policy.

## Introduction

There is a growing interest in characterizing genome-wide variability and genetic structure at small or regional geographic scales in different human populations. Several national European examples include extensive GWAS studies performed in Britain (Leslie *et al*., 2015), France (Karakachoff *et al*., 2015), Finland (Kerminen *et al*., 2017), Italy (Fiorito *et al*., 2016), Ireland (Byrne *et al*., 2018), Scotland (Gilbert *et al*., 2019), and Spain (Bycroft *et al*., 2019). This renewed interest in fine-scale genetic structure is essential for controlling genetic ancestry in common variant association studies (Mathieson & McVean, 2012), as well as understanding rare variant association studies, since rare variants are often specific to certain geographically clustered populations (Genomes Project *et al*., 2015). In addition, fine-scale genetic structure can clarify the relationships between closely related populations and uncover recent historical events. However, there is a scarcity of studies performed with sequencing data, since most of these population studies have been carried out using genome-wide genotypic data; an early notable exception being the Genome of the Netherlands Project (Genome of the Netherlands, 2014). Large-scale sequencing is far more precise and informative in revealing genetic structure, and it also allows for novel variants to be discovered. In the Iberian Peninsula, the shortage of large-scale genomic studies is even more noticeable, with, to our knowledge, only three studies published, one using Whole Exome Sequencing (WES) data (Dopazo *et al*., 2016), and the other two using Whole-Genome Sequencing (WGS), one of them focusing in the northeastern Iberian region of Catalonia (Valls-Margarit *et al*., 2022), and the other primarily addressing signals of natural selection (Garcia-Calleja *et al*., 2025).

Galicia is a region occupying the northwestern corner of the Iberian Peninsula that possesses several unique characteristics. Its geographical isolation from the rest of the peninsula has, to a certain degree, maintained its identity and distinctive culture, including its own language. Besides, at the genomic level, Galicia seems to show a special differentiation, with examples of possible founder effects related to several recessive Mendelian diseases such as ichthyosis (Esperón-Moldes *et al*., 2019) or Wilson disease (Brage *et al*., 2007), as well as exhibiting signs of local genetic inbreeding (Esperón-Moldes *et al*., 2019) or isolation and low genetic diversity (Salas *et al*., 1998). Its peripheral location in Europe and its relative isolation make Galicia particularly intriguing for studying population genetic structure. Recent data on the Spanish population by Bycroft et al. (2019) has reported extraordinary fine-scale population substructure within Galicia, albeit using genotypic data (∼693K SNPs) and exhibiting a geographically biased sampling towards the southwestern region of the Pontevedra province, where the great majority of the Galician samples were recruited.

Previous studies have reported a significant North African genetic contribution to Iberian populations, particularly in Galicia, believed to stem primarily from Arab-Berber migrations during the Muslim conquest of the Iberian Peninsula in 711 CE. Islamic rule lasted in various forms until the fall of Granada (in Andalusia, in the southern part of the peninsula) in 1492. The Kingdom of Asturias, which encompassed present-day Galicia, served as a key Christian stronghold against Muslim expansion. Over time, this resistance evolved into the *Reconquista*, a prolonged series of military campaigns aimed at reclaiming territories under Islamic rule. By the mid-8th century, Galicia had largely returned to Christian control. Therefore, Galicia played a pivotal role in the broader *Reconquista* efforts as this region served as a refuge for Christian elites and populations, and from there, the first resistance movements began. While the direct impact of Muslim rule in northern Iberia, including Galicia, was relatively limited compared to other regions, genetic studies suggest a substantial proportion of Galician ancestry originated from North Africa. The availability of WGS data now provides an unprecedented opportunity to investigate this ancestry with higher genetic resolution than previous studies.

In this study, we generated WGS data from 94 individuals of Galician origin, a sample size comparable to that of other WGS studies conducted in Iberia and Europe. The availability of high-resolution genomic data enabled us to investigate inbreeding patterns and explore the extent of African and Middle Eastern admixture in Galicia. We analyzed genetic variation in the region at the highest level of resolution to build upon previous remarkable findings suggesting the presence of ultrafine population structure in Galicia (Bycroft *et al*., 2019). To further evaluate these findings, we additionally genotyped samples from across Iberia (*n* = 453), closely replicating the sampling strategy of Bycroft et al. (2019) to critically assess their reported results in the Galician population and, more broadly, in Spain. Finally, we conducted the first investigation into the biomedical implications of fine-scale population structure within Galicia, analyzing micro-spatial patterns of polygenic risk for various common diseases.

## Material and Methods

### Sampling

We collected 94 DNA samples (hereafter referred to as the GALOMICS [GAL] dataset) from donors representing the entire territory of Galicia (northwest Spain), ensuring maximum geographic homogeneity in the sampling process. Of these, 91 donors passed all the filters for subsequent analysis (see below). The average distance between the donors’ birthplaces and those of their four grandparents was 6.6 km (SD: 15.8), with a median of 0 km (IQR: 0–5.5 km). The average distance among the birthplaces of the four grandparents was 5.4 km (SD: 10.7), also with a median of 0 km (IQR: 0–5.2 km). No missing data were recorded regarding the biogeographical origins of the donors or their grandparents. The sampling primarily represents a rural population, defined as areas with fewer than 10,000 inhabitants (according to official statistics: https://www.ige.gal/). Specifically, the median population size of the donors’ birthplaces was 9,080 inhabitants, with 59.3% of donors born in rural areas as defined above. Additionally, the median population size of the birthplaces of the donors’ grandparents was 5,168 inhabitants. In total, 63.7% of donors have all four grandparents born in rural areas, 75.8% have at least two grandparents from rural areas, and only 20.8% have all four grandparents born in urban areas. **Figure S1 – Supplementary File**. By province, the samples were distributed as follows: 39.6% from A Coruña, 20.9% from Ourense, 23.1% from Lugo, and 16.5% from Pontevedra.

Saliva samples were collected using Oragene OG-500 devices (DNA Genotek), and DNA was then isolated using the prepIT L2P kit from DNA Genotek, following the standard protocol. Written informed consent was obtained from all Galician donors before the research. The rights of participants were safeguarded during the research, and their identities were protected. The study complies with all applicable Spanish regulations, including the Biomedical Research Act (14/2007-3 of July), the Autonomy of the Patient Act (41/2002), Decree SAS/3470/2009 for observational studies, and the Data Protection Act (15/1999). The experimental protocol was approved by the Galician Ethics Committee (registration code: 2017/399).

In addition to performing WGS on the GAL dataset, we genotyped DNA samples from the Spanish National DNA Bank (https://www.bancoadn.org/en/presentation.html; hereafter referred to as the NDNAB dataset). These samples were collected from 453 donors representing various Spanish provinces across different autochthonous regions, including all four provinces of Galicia. Donors in this dataset were selected to ensure broad regional representation in the Spanish territory, somewhat comparable to the sampling strategy used by Bycroft et al. (2019); samples from the Spanish National DNA Bank were, in fact, also used by these authors.

### Reference Populations

We used three different repositories for WGS data of reference populations. From the 1000 Genomes Project (1000G; https://www.internationalgenome.org), we accessed data representing major European, Asian, and African populations. Variant files were initially downloaded from the 1000G public FTP site (ftp://ftp.1000genomes.ebi.ac.uk/vol1/ftp/release/20110521/). For the bioinformatic processing of the 1000 Genomes data (http://ftp.1000genomes.ebi.ac.uk/vol1/ftp/phase1/data), we leveraged previous bioinformatic developments (Amigo *et al*., 2011; Amigo *et al*., 2008). From the Human Genome Diversity Project (HGDP), we used data from seven datasets representing additional whole genomes from Europe, North Africa, and the Middle East (Cann *et al*., 2002; Cavalli-Sforza, 2005). The downloaded data from the 1000G contained 78,090,318 SNPs, while the HGDP dataset included 64,214,739 SNPs. The combined overlapping data from all these population datasets (GAL + 1000G + HGDP) consisted of 11,085,935 SNPs. To better understand the African contribution to the Galician genomes, we enriched the admixture analyses by incorporating North African and Middle East WGS data from Serra-Vidal et al. (2019). A summary of the datasets used in this study is provided in **Table 1**, and **Figure 1A** shows a map of the datasets included.

**Figure 1.**
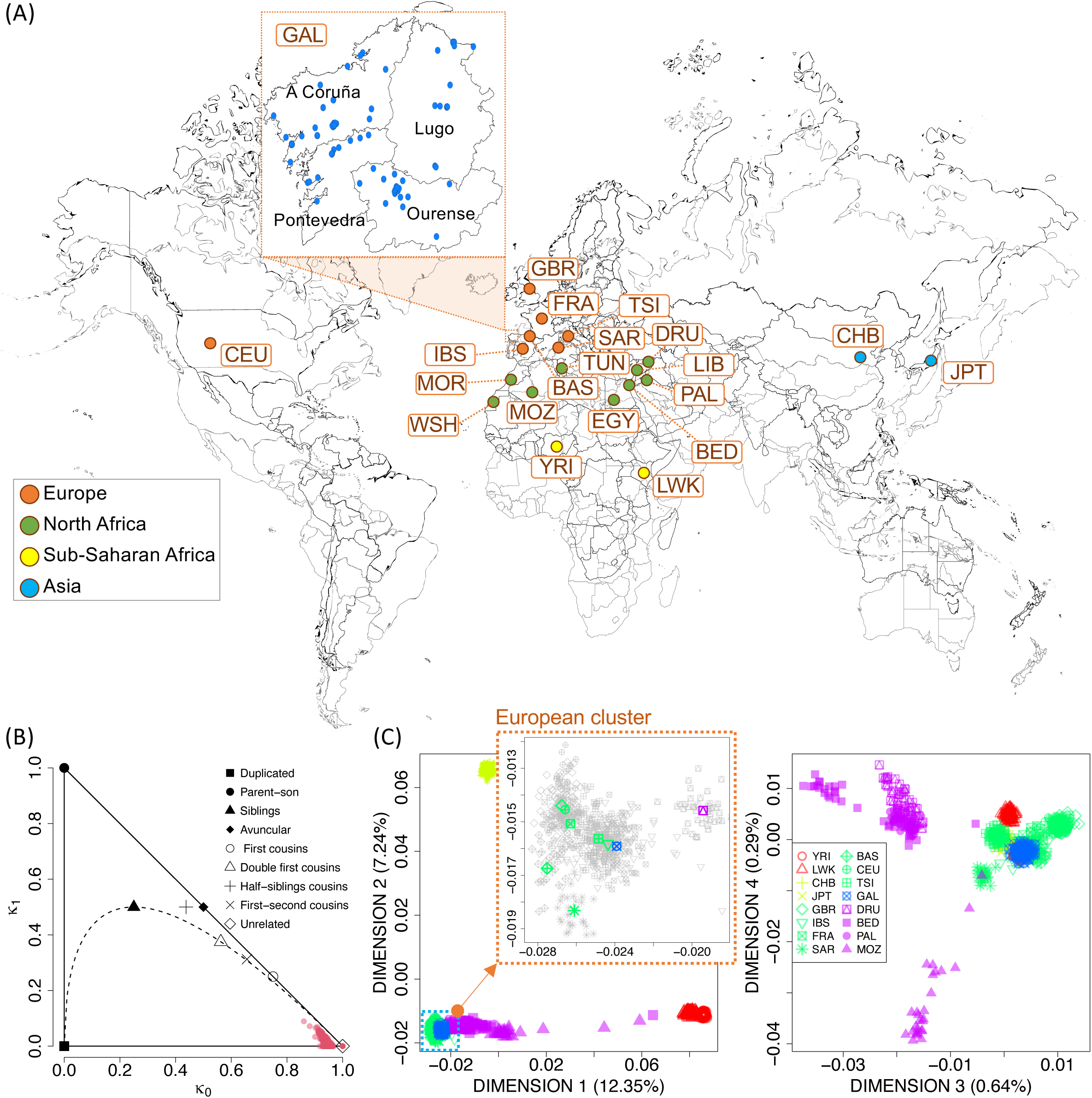
(A) Map illustrating the datasets used in this study, with a zoomed-in section providing details on the Galician (GAL) dataset sampling locations (indicating also the four provinces of the region: A Coruña, Lugo, Ourense and Pontevedra). (B) Kinship analysis of the GAL dataset, showing that after filtering out closely related individuals (removed from the plot), the remaining samples used in this study exhibit no close familial relationships. (C) MDS analysis comparing the GAL dataset to major continental populations. The left MDS plot includes a zoomed-in view of the European genomes, highlighting the centroids of the population datasets. See **Table 1** for population code details.

**Table 1.** Population WGS datasets used in the present study.

| **Population/Ethnicity** | **Country/Region** | **Acronym** | ***Continental region*** | ***N*** | **Source** |
| --- | --- | --- | --- | --- | --- |
| Han Chinese in Beijing | China | CHB | Asia | 103 | 1000G |
| Japanese in Tokyo | Japan | JPT | Asia | 104 | 1000G |
| Basque | Spain | BAS | Europe | 23 | HGDP |
| British | England/Scotland | GBR | Europe | 91 | 1000G |
| French | France | FRA | Europe | 28 | HGDP |
| Galicians (present study) | Spain | GAL | Europe | 91 | 1000G |
| Iberian Population | Spain | IBS | Europe | 107 | 1000G |
| Sardinian | Italy | SAR | Europe | 28 | HGDP |
| Toscani | Italy | TSI | Europe | 107 | 1000G |
| Utah Residents (CEPH) | USA | CEU | Europe | 99 | 1000G |
| Bedouin | Middle East | BED | Middle East | 46 | HGDP |
| Druze | Middle East | DRU | Middle East | 42 | HGDP |
| Palestinian | Middle East | PAL | Middle East | 46 | HGDP |
| Berbers/Moroccans | Morocco | MOR | North Africa | 4 | Serra-Vidal et al. (2019) |
| Chenini Berbers/Tunisians | Tunisia | TUN | North Africa | 4 | Serra-Vidal et al. (2019) |
| Egyptians | Egypt | EGY | North Africa | 2 | Serra-Vidal et al. (2019) |
| Libyans | Libya | LIB | North Africa | 2 | Serra-Vidal et al. (2019) |
| Mozabite | Algeria | MOZ | North Africa | 27 | HGDP |
| West-Saharans | West Sahara | WSH | North Africa | 1 | Serra-Vidal et al. (2019) |
| Zenata Berbers/Algerians | Algeria | ALG | North Africa | 4 | Serra-Vidal et al. (2019) |
| Luhya in Webuye | Kenya | LWK | Sub-Saharan Africa | 99 | 1000G |
| Yoruba in Ibadan | Nigeria | YRI | Sub-Saharan Africa | 108 | 1000G |

### Whole-Genome Sequencing

Sequencing was conducted by BGI using the following protocol. Genomic DNA samples were randomly fragmented using Covaris technology, producing fragments of approximately 350bp. After fragment selection, DNA fragments underwent end repair, followed by the addition of an “A” base to the 3’ end of each strand. Adapters were ligated to both ends of the end-repaired/dA-tailed DNA fragments. This was followed by ligation-mediated PCR (LMPCR) amplification, single-strand separation, and cyclization. Rolling circle amplification (RCA) was then performed to create DNA Nanoballs (DNBs). The qualified DNBs were loaded onto patterned nanoarrays, and paired-end reads were generated using the BGISEQ-500 platform. High-throughput sequencing was performed for each library to ensure that the average sequencing coverage requirement for each sample was met. The sequencing-derived raw image files were processed using the BGISEQ-500 base-calling software with default parameters to generate the sequence data as paired-end reads. This sequence data, referred to as “raw data,” was stored in FASTQ format.

### Genotyping of the Spanish DNA National Bank

Samples of the NDNAB dataset were genotyped using the Axiom® Genome-Wide Human Origins 1 Array at the Centro Nacional de Genotipado (CEGEN) in Santiago de Compostela, Spain. Following sample processing, genotype calls and quality control metrics were generated using the Affymetrix Genotyping Console™ (GTC) 4.1.2 software, adhering to standard commercial guidelines. As part of this process, sample dish QC (DQC) values were calculated, and any sample with a DQC value below the default threshold of 0.82 was excluded from the study. Genotyping was performed on the remaining high-quality samples using the “AxiomGT1_all” algorithm, incorporating all SNPs available on the array. In total, 599,424 SNPs were successfully retrieved from the NDNAB dataset.

### Bioinformatics Analysis

Quality control (QC) of DNA FastQ was performed using *FastQC* v0.11.9 [1] and *MultiQC* v1.10 (Ewels *et al*., 2016). They were mapped to Human Genome v37 obtained from (https://console.cloud.google.com/storage/browser/gcp-public-data--broad-references/hg19/v0) using BWA 0.7.17-r1188 (Li, 2013; Li & Durbin, 2010). PCR duplicates were marked using *Picard* 2.27.0 (https://broadinstitute.github.io/picard/). Variants were discovered using *Google Deepvariant* 1.5.0 (Poplin *et al*., 2018) and *GLnexus* (Yun *et al*., 2021). The Combined Annotation Dependent Depletion (CADD) *GRCh38*-v1.6 dataset (Rentzsch *et al*., 2021) was used to obtain predictive information quantifying the deleteriousness of SNPs and small insertion-deletion variants (indels) across the human genome. This database integrates a wide range of annotations to assess the potential impact of variants, encompassing both coding and non-coding regions. CADD provides two main types of scores: raw scores, which reflect the unscaled deleteriousness of a variant based on the CADD model, and PHRED-scaled scores (used in the present study), which are log-transformed and adjusted for easier interpretation. Additionally, the dataset includes information on the functional classification of variants, such as whether they are non-synonymous, synonymous, stop-gained/lost, intronic, or fall into other categories such as regulatory regions or splice sites.

### Analysis of Uniparental Markers

Information on mitochondrial DNA (mtDNA) and Y-chromosome variation in GAL was extracted from the WGS data. MtDNA variation was contrasted against the rCRS, and haplotypes were classified into haplogroups using *HaploGrep* software (Weissensteiner *et al*., 2016). To infer Y-chromosome haplogroups from WGS data, we used the Python-based tool *Yleaf* (Ralf *et al*., 2018).

### Biogeographical analysis

To investigate clusters of genome variation, we first carried out Multidimensional Scaling (MDS) analysis using pairwise Identity-By-State values of the GALOMICA sample and the reference populations. MDS was performed using the function *cmdscale* (library *stats*) from R (http://www.r-project.org). Average Identity-By-State distances were computed as the average of the matrix of identity-by-state distances computed between individual genomes against all the individuals from each reference population.

### Admixture Analysis

Maximum likelihood estimation of individual ancestries from multi-locus SNP data was undertaken using ADMIXTURE (Alexander *et al*., 2009); this software uses a maximum likelihood estimation of individual ancestries from multi-locus SNP data. For the analysis, we used the following filtering criteria: MAF > 0.05 and LD pruning with a window size of 100bp, a step size of 10, and *r*^2^ > 0.5, following the same approach used to compute inbreeding coefficients (Chang *et al*., 2015), as detailed below. We incorporated various sources of worldwide populations from the 1000G and the HGDP. First, we run ADMIXTURE in an unsupervised mode to gain a general view of the potential demographic influence of Africa on the GAL dataset. Shared ancestry between Mediterranean populations from Europe and Africa could lead to an overestimation of North African and Middle Eastern ancestry in Galicians. To address this, we also used supervised ADMIXTURE analysis using a minimal set of reference samples representing the most likely African-origin surrogates for Galician ancestry with the datasets available. In this supervised analysis, we included only genomes in which >80% or 90% of their genetic makeup was assigned to the respective North African or Middle Eastern components, thereby minimizing the impact of shared genetic influence with Europeans.

We computed *f_3_*–statistics to assess whether the genetic relationships among the three populations could be explained with or without admixture. Specifically, we used the three-population test in the form *f_3_*(X_EUR_,X_NAF/SSH_;GAL), where (X_EUR_) represents a particular European population subset, X_NAF/SSH_ denotes a specific North African or Sub-Saharan African population sample, and GAL corresponds to the Galician dataset. This test evaluates whether the genetic makeup of Galician genomes can be attributed to admixture between European and North African populations. A significantly negative *f*_₃_-value (typically with Z-score ≤ –3) is strong evidence that GAL has experienced admixture between populations related to X_EUR_ and X_NAF/SSH_. A positive *f*_₃_-value or non-significant Z-score suggests no detectable admixture, or that the populations chosen (X_EUR_, X_NAF/SSH_) do not adequately model the admixture sources. To ensure the robustness of our conclusions, we tested multiple combinations of European and African datasets, capturing the genetic diversity within these regions.

### Dating Admixture

We used ALDER (Loh *et al*., 2013) to estimate the admixture timing of a single source population from North Africa (represented by Mozabites) and Middle East (represented by Bedouins) into our recipient population, Galicians (GAL), by analyzing the LD decay within the recipient group: When a population receives genetic input from an external group, the introduced alleles initially exhibit long-range LD, which gradually decays over generations due to recombination. Input files for ALDER were prepared using EIGENSOFT (Patterson *et al*., 2006). The ALDER framework assumes random mating in the target population following the admixture event and is optimized for detecting relatively recent admixture, whether arising from a single pulse or multiple waves of gene flow (Moorjani *et al*., 2011).

We used fastGLOBETROTTER following the author’s recommendations (Wangkumhang *et al*., 2022) to independently infer and date admixture events. As a first step, we used CHROMOPAINTER (Lawson *et al*., 2012) to generate haplotype-sharing profiles by modeling each individual’s genome as a mosaic of segments copied from a set of reference populations. These haplotype profiles were then analyzed by fastGLOBETROTTER, which examines the distribution and decay of shared haplotype segments across the genome, a signal shaped by historical recombination. By assessing how ancestry contributions from different source populations vary with genetic distance, fastGLOBETROTTER infers the number, timing, and ancestral composition of admixture events. Unlike methods that rely solely on allele frequency differences, fastGLOBETROTTER derives its high resolution by using haplotype structure, which enables it to detect even subtle or ancient admixture signals with greater sensitivity. In this study, we applied fastGLOBETROTTER specifically to detect and characterize admixture signals involving North African and/or Middle Eastern ancestral sources, similar to our use of ALDER. To simplify model complexity, and given the limited availability of WGS reference datasets, our model included representatives for four broad ancestry components: sub-Saharan variation (Yoruba from the 1000 Genome Project), European ancestry (CEU from the 1000G), North African (Mozabites from the HGDP), and Middle Eastern (Bedouin from the HGDP).

To reduce the confounding effects of European-like variation in African populations, we excluded Mozabites and Bedouin individuals with <80% African-like ancestry. Model comparison indicated that a two-date admixture model did not improve the fit relative to a single-date model. Therefore, we report results from the simpler, single-date admixture scenario, which we consider the most parsimonious representation of the underlying demographic history. We also report 95% confidence intervals for admixture dates based on 200 bootstrap resamples, providing robust statistical uncertainty.

Despite the low-resolution nature of the NDNAD dataset and its limited sample size, we also aimed to explore temporal signals across different geographic regions, to evaluate whether there is evidence supporting differential entry dates of North African and Middle Eastern ancestry into the Iberian Peninsula. In particular, we focused on possible differences between the northern regions and the central/southern parts of the Peninsula. This question arises from the widespread assumption that the Arab rule, beginning in 711 CE and followed by the gradual Christian *Reconquista*, was the primary source of North African ancestry in Iberia. However, there is limited historical support for a strong Arab-Berber presence in the far North of the Peninsula, where the Muslim political and military influence was more restricted and often transient. Consequently, while the Islamic conquest could plausibly account for much of the North African genetic signal in central and southern Iberia, the presence of this component in the North may reflect alternative demographic processes and earlier or later migratory events, potentially including Roman-era, pre-Islamic, or post-*Reconquista* population movements.

fastGLOBETROTTER also provides estimates of ancestry proportions from the reference populations considered (e.g. Mozabites and Bedouins), which are expected to be consistent with the proportions inferred using ADMIXTURE.

We used the latest estimate of generation time by Wang et al. (2023), which provides one of the most comprehensive and robust empirical estimates of human generation intervals, based on a genomic approach using *de novo* mutations (DNMs). According to these authors, for a typical European population over the past 40,000 years, the sex-averaged generation time is estimated at 26.1 years.

### Analysis of FineSTRUCTURE

Clusters of individuals in the Galician dataset were inferred using fineSTRUCTURE (Lawson et al., 2012) following the author’s recommendations. The number of clusters is not determined *a priori*; it is just the number estimated under the underlying probability model. We used the CHROMOPAINTER package (Lawson et al., 2012) to estimate the coancestry matrix derived from the clusters estimated by the software and to build a phylogenetic tree of clusters. FineSTRUCTURE was first run on the GAL dataset to investigate population clusters within the Galician territory. To extend the analysis to a national scale and contextualize Galicia within Spain, we analyzed the clustering pattern in the NDNAB dataset.

Principal Component Analysis (PCA) analyses based on the coancestry matrix were generated using R scripts provided by the authors (available at http://www.paintmychromosomes.com/finestructure/).

### Analysis of Runs of Homozygosity and Inbreeding

Runs of homozygosity (ROH) and inbreeding were assessed in GAL and other reference populations using WGS data analyzed in PLINK (Chang *et al*., 2015). ROH segments were identified following previous protocols (Serra-Vidal *et al*., 2019), applying sliding windows of 50 SNPs across the genome, defining segments containing a minimum of 50 SNPs within 100 kbp, allowing gaps of up to 2000 kbp between consecutive SNPs, with no more than one heterozygous site and five missing sites. To evaluate differences between GAL and other populations, Wilcoxon tests were computed on multiple ROH metrics: total counts of short and long ROH segments (*nROH_S_* and *nROH_L_*; respectively), cumulative length of short and long ROH segments (*sROH_S_*and *sROH_L_, respectively*), and average length of short and long ROH segments (*mROH_S_* and *mROH_L_, respectively*); here, short segments are defined as those ≤1.5 Mb and long segments as those >1.5 Mb. We also computed *F_ROH_,* which estimates the total inbreeding coefficient (estimator of F_IT_ of Wright’s fixation index (Hartl & Clark, 2007)) and is defined as the sum of ROH segments >1.5 Mb divided by the total length of the genome in Mb (in our study, 2,795 megabases [Mb]).

We additionally computed four single-point estimators of inbreeding coefficients based on allele frequencies. The coefficient *F_HET_* measures the average SNP homozygosity within an individual relative to the expected homozygosity of alleles randomly drawn from the population (genome-wide homozygous excess due to inbreeding). *F_hat1_* represents the variance-standardized relationship minus 1 (equivalent to the diagonal of the covariance matrix used for PCA). *F_hat2_*is based on the homozygous excess. The estimator *F_HET_* is analogous to the *F_HAT2_* estimator, both measuring the excess of homozygosity, but they are not identical; while *F_HET_* is a ratio of sums, *F_HAT2_* is a sum of ratios (Gazal et al., 2014). *F_hat3_* is based on the correlation between uniting gametes; it uses the initial definition of the inbreeding coefficient proposed originally by Sewall Wright in 1922. PLINK v1.9 (Chang *et al*., 2015) software was used to curate the data and perform most of the analyses. R statistical software (The R Core Team, 2019) was used for the computation of identity-of-state values used in the MDS analysis, ROH, and F inbreeding metrics, and graphic representations.

### Exploring Variants of Biomedical Interest

We are interested in exploring genomic variation in Galicia on a WGS scale, particularly its biomedical relevance compared to the broader Iberian population. This could help anticipate the potential benefits of conducting large-scale whole-genome projects at a regional level, rather than focusing solely on, e.g. exome variation, which, while more cost-effective, may not capture the full spectrum of relevant genetic diversity.

First, we conducted a single-point allele association test, comparing single-nucleotide polymorphism (SNP) variants detected in GAL with those in IBS (*n* = 107) and a merged European dataset (CEU + GBR + IBS + TSI; *n* = 404). This approach mimics a GWA’s study, where typically a cohort of cases (such as GAL) is compared to a cohort of controls (such as IBS) (Salas *et al*., 2017). For this analysis, we included only variants with a minor allele frequency (MAF) above 0.05 in cases and controls and a Hardy-Weinberg equilibrium *P*-value > 0.001 for GAL, IBS, and the European merged dataset.

Second, to investigate the impact of accumulated pathogenic variants in genes that could increase susceptibility to common diseases in Galicia compared to other Iberian/European populations, we performed a statistical association test using the collapsing method known as the Optimized Sequence Kernel Association Test (SKAT-O) (Lee *et al*., 2012). This method combines a set of generalized SKAT tests with varying proportions of SKAT and Burden. We incorporated annotated information for each variant obtained from the Combined Annotation Dependent Depletion (CADD) v1.6. database (Rentzsch *et al*., 2021). We undertook this gene-based statistical analysis using the *SKAT* R library (Lee & Miropolsky, 2023) and considering all variants within genes, aiming to capture rare variations that could be of biomedical interest and might go unnoticed in single-point association tests. Genes with at least two variants were considered for these analyses. A nominal significance level was set to 0.05. To monitor multiple tests, we used the Bonferroni correction.

The top SNP and gene candidates from these analyses were further investigated for their association with medical conditions. For this, we gathered information from SNPedia, (downloaded from https://www.snpedia.com/index.php/Category:Is_a_snp [for SNP variation] and https://www.snpedia.com/index.php/Category:Is_a_gene [for genes], updated in October 2023) and classified it into ten macro-areas (October 2024). The SNPedia contains information on 111,702 genetic variants and 2,164 genes (https://www.snpedia.com/index.php/SNPedia:FAQ#How_many_SNPs_are_in_SN Pedia.3F).

Third, given the growing interest in predicting genetic predisposition to diseases and traits within the biomedical community, we analyzed potential regional differences in Polygenic Risk Score (PRS) predictions, comparing PRS values between the GAL and IBS populations. PRSs estimate disease risk based on the cumulative effect of multiple genetic variants, typically inferred from large-scale GWAs. To streamline our analysis, we selected PRSs for common diseases with significant public health relevance, including five genome-wide PRSs from Khera et al. (2018): *i*) Coronary Artery Disease (PRS_CAD_; PGS000013): 6,630,150 polymorphisms; 6,476,712 retained after merging GAL and IBS; *ii*) Atrial Fibrillation (PRS_AF_; PGS000016): 6,730,541 polymorphisms; 6,562,700 retained; *iii*) Type 2 Diabetes (PRS_T2D_; PGS000014): 6,917,436 polymorphisms; 6,715,496 retained; *iv*) Inflammatory Bowel Disease (PRS_IBD_; PGS000017): 6,907,112 polymorphisms; 6,712,364 retained; and *v*) Breast Cancer (PRS_BC1_; PGS000015): 5,218 polymorphisms; 4,532 retained. Additionally, we included: *i*) a second Breast Cancer PRS metric (PRS_BC2_; PGS000004), implemented in the CanRisk calculator (https://www.canrisk.org: 313 polymorphisms, 286 retained; *ii*) Ovarian Cancer (PRS_OCAC_; PGS003394): 36 polymorphisms; 36 retained (Lee *et al*., 2019), also implemented in CanRisk, *iii*) Acute Lower Respiratory Infection (PRS_ALRI_; PGS000925): 402 polymorphisms, 346 retained; the only infectious disease PRS indexed in the PGS Catalog to date, included due to various EWAS carried out on infectious diseases in the Galician population to date (Camino-Mera *et al*., 2024; Pardo-Seco *et al*., 2024; Salas *et al*., 2018), iv) Schizophrenia (PRS_SCH_; PGS000133): 604,645; 602,522 retained (Zheutlin *et al*., 2019), *v*) Autism Spectrum Disorder (PRS_ASD_; PGS000327): 35,087 polymorphisms; 34,362 retained (Grove *et al*., 2019), and Alzheimer’s Disease (PRS_ALZ_; PGS004146): 915,771 polymorphisms; 914,409 retained (Monti *et al*., 2024). This selection allows for the exploration of PRSs composed of a wide range of polymorphisms, from a few dozen to several million, capturing different levels of genome-wide architectures. Notably, ∼98% of the variants included in these PRSs are in non-coding regions of the genome, which would not be detected in EWS studies. While these non-coding variants do not alter protein sequences, they can influence gene expression and contribute to disease susceptibility. PRS calculations were performed using PLINK (Chang *et al*., 2015), and values were standardized to a unit standard deviation (SD) within the GAL + IBS pooled dataset.

In addition, we have developed a web tool (https://galomica.genpob.eu/) that enables users to search for specific variants in the GALOMICS whole genome database, which contains information on over 18 million biallelic variants. In the “Search” tab, users can query genetic variants by two main criteria: by rsID (e.g., rs1805005) or by a combination of chromosome and position (e.g., 16: 89919436). The following visualizations are provided: *i*) A geometric map displaying the coordinates of the samples, including layers for individuals as well as their parents and grandparents., *ii*) Various charts indicating the samples in GAL that carry the reference allele *vs*. those with the alternative allele, showing the genotypic composition of individuals, *iii*) A representation of allele variant frequencies across different populations worldwide, and *iv*) Biomedical data of interest. For graphical representations of allele frequencies, data is sourced from external databases, including: *i*) Ensembl (http://rest.ensembl.org); Entrez: ClinGen: http://reg.clinicalgenome.org/redmine/projects/registry/genboree_registry/by_caid? caid=; UniProtKB: http://www.uniprot.org/uniprot/; OMIM: http://www.ncbi.nlm.nih.gov/omim/; dbSNP: http://www.ncbi.nlm.nih.gov/snp/. The data obtained from these sources is structured in JSON format. Additionally, a ClinVar Data Table displays the identifiers assigned to the queried variant by ClinGen, UniProtKB, OMIM, and dbSNP. The table includes details such as the cDNA Change (alteration at the complementary DNA level), the variant’s position in the hg38 version of the human genome, the associated Trait Name, the Trait Reference, and the Protein Change (protein alterations caused by the variant).

### Visualization of Genetic Data in Geographic Maps

To visualize the biogeographical distribution of genetic cluster frequencies, we first calculated individual cluster frequencies at each sampling location. These frequencies were then spatially represented on geographic maps using SAGA v. 9.6.1 (http://www.saga-gis.org/) (Conrad *et al*., 2015) and the ordinary Kriging interpolation method. For the GAL samples, we had detailed information on the specific birthplaces of donors and their grandparents. In contrast, for the NDNAB samples, data resolution was limited to the provincial level. Therefore, for practical purposes, each data point was assigned to the capital city of the corresponding province. Consequently, each point in this case represents all individuals from that province, sharing the same geographic coordinates.

## Results

### Overview of Galician WGS Variation

We first removed non-autosomal variants, indels, and variants with a genotyping rate below 99%. After applying additional filters (e.g., removal of triallelic SNPs and indels), the dataset was reduced to over 17.2 million variants, which were used for downstream analyses (unless additional filtering was required, e.g., for Hardy-Weinberg disequilibrium, linkage disequilibrium [LD]). Approximately 36.4% of the variants were located within intronic regions, while 32.3% were in intergenic regions. Additionally, 15.2% of the variants were identified in regulatory or splice site regions, and 0.8% were in exons. Among the variants located in exons, >35K were non-synonymous, >29K were synonymous, and >79K were found in 3’/5’-untranslated regions (UTR). The remaining 500 variants were classified as either stop-gained or stop-lost.

We first investigated the familial relationship between donors in GALOMICS as done previously (Camino-Mera *et al*., 2024; Pardo-Seco *et al*., 2016). Of the initial 94 WGS, three were excluded due to close relationships (**Figure 1B**) or their behavior as outliers in the MDS analysis (see below).

Finally, we generated a web tool, GALOMICS (https://galomics.genpob.eu), to facilitate the search for allele variants in the GAL database. The tool has been designed to be scalable, anticipating the integration of new variants from other datasets. Many of these datasets represent disease conditions in Galician donors, some of which have already been generated, e.g. (Butler-Laporte *et al*., 2022; Camino-Mera *et al*., 2024; Pardo-Seco *et al*., 2024; Salas *et al*., 2018; Salas *et al*., 2017).

### Biogeography and Admixture Analysis of Galician Genomes

As expected, the MDS plot (**Figure 1C**) of worldwide population genomic datasets shows most samples clustering around the three main continental groups: sub-Saharan Africa, Asia, and Europe. The first dimension (accounting for 12.35% of the variation) primarily separates the sub-Saharan group (LWK and YOR) from the rest, while Dimension 2 (7.24% of the variation) distinguishes Central Asians (CHB and JPT) from Europeans (represented by BAS, CEU, FRA, GAL, GBR, IBS, SAR, and TSI) and sub-Saharan Africa. The North African datasets (BED, DRU, MOZ, and PAL) are more closely related to the European cluster, slightly shifted towards the sub-Saharan vertex, with a few exceptional profiles very closely related to sub-Saharans. In a close-up view of the European/North African cluster (**Figure 1C**; left), Galician genomic profiles are embedded within the European cluster but are not positioned at its core. Instead, they show a discrete shift towards the North African pole. Dimension 3 (0.64%; **Figure 1C**; right) separates North Africans from Europeans, with Bedouins (BED) at the opposite pole from Europeans, while Dimension 4 (0.29%) separates North Africans and Europeans from Mozabites (MOZ; **Figure 1C**; right). These two dimensions highlight Bedouins and Mozabites at the extremes of North African variation, with the rest of the North African samples showing more genetic affinity to Europeans.

We used the unsupervised clustering algorithm of ADMIXTURE to estimate ancestry proportions in Galician genomes, which are influenced by gene flow between African and European populations (**Figure 2A**). Our analysis utilized population datasets representing major continental and potentially ancestral groups, including sub-Saharan Africa, North Africa, Asian, and Europe. We enriched this analysis with other North African samples analyzed in Serra-Vidal et al. (2019). We explored values of *K* ranging from 2 to 10 ancestral contributors, but we only discuss results for *K* = 3 and 5, because these are relevant in the present demographic context. At *K* = 3, ADMIXTURE identifies ancestry components originating from three main geographic locations, sub-Saharan Africa, Central Asia, and North Africa/Europe. When *K* = 4, the model consistently distinguishes the North African component from the primary European one, suggesting a shared ancestry between Europe and North Africa (**Figure 2A**; left). At *K* = 5, which corresponds to the model with the lowest cross-validation value in the admixture analysis, the algorithm further divides the North African/Middle East component into two subcomponents. One of these subcomponents mainly characterizes North African Mozabites (hereafter referred to as ‘Mozabites North African’-like ancestry), while the other reflects the variation mainly observed in the Middle East (represented by Palestinian, Bedouin and Druze) and the other North African datasets (hereafter referred as to ‘non-Mozabites North African’-like ancestry). All the datasets in North Africa and the Middle East have, however, variable proportions of these two ancestries. Moreover, these two North African ancestries are also present in other European datasets FRA, SAR, TSI, IBS, and GAL, while the ‘non-Mozabite North African’-like ancestry is absent in the BAS, and both North African ancestries are absent in CEU and GBR.

**Figure 2.**
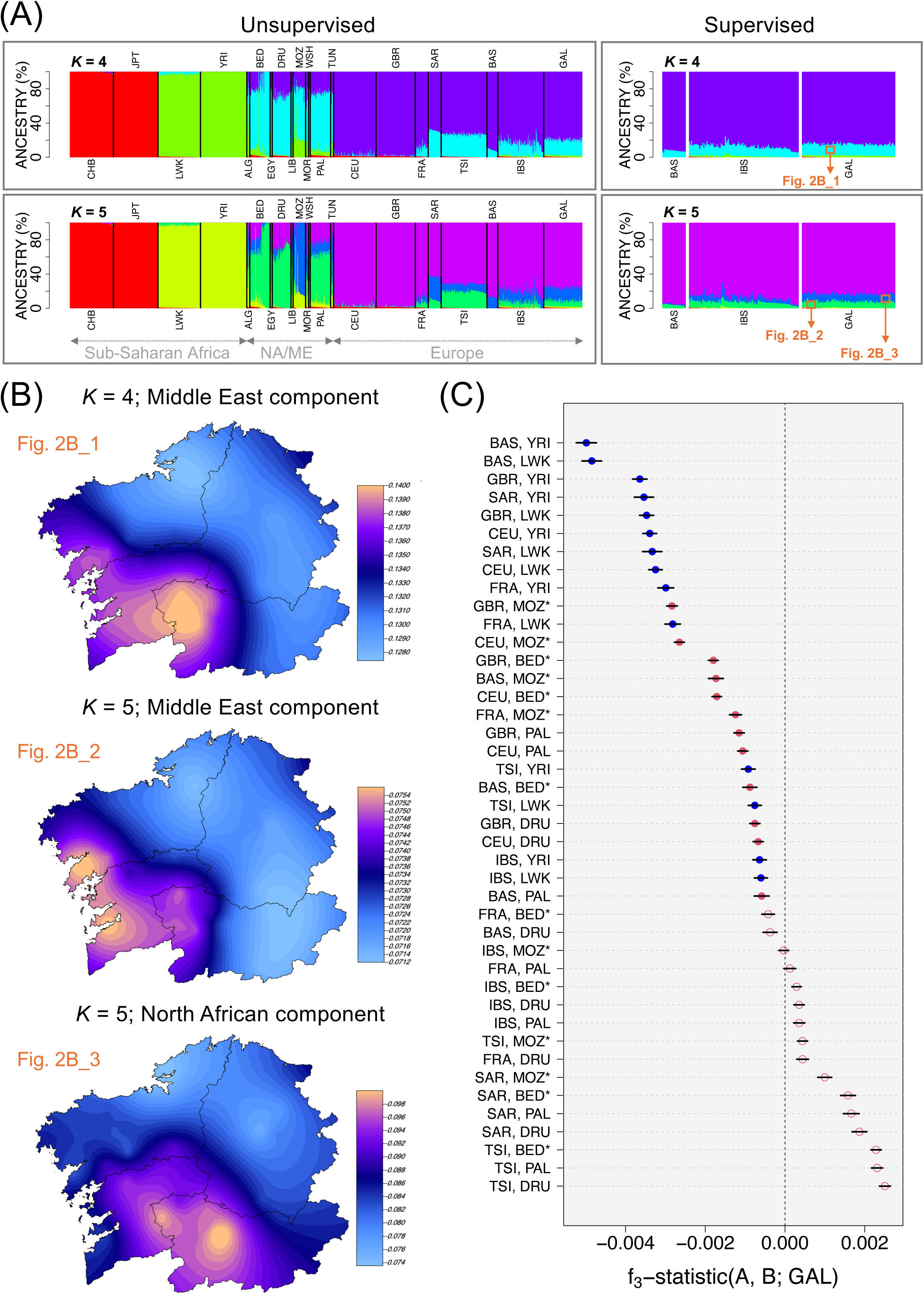
(A) Bar plot depicting individual ancestry proportions estimated using ADMIXTURE with both unsupervised (left) and supervised (right) clustering approaches, considering K values from 4 to 5. (B) Interpolated frequency map of the Middle East and North African component among GAL donors across the Galician region, based on the ancestry component identified in (A) (see indications for Fig. 2_1; Fig.2_2, and Fig.2_3). (C) *f*_₃_-statistics in the form *f*_₃_(X_EUR_, X_NAF/SSH_; GAL), measuring shared genetic drift. Each dot represents an f_₃_ value calculated for a pair of source populations with GAL as the target population. Blue dots correspond to combinations involving sub-Saharan African populations, red dots indicate combinations with North African or Middle East populations, and filled red dots highlight statistically significant results (Z-score ≥ 3), suggestive of admixture events contributing to the genetic makeup of the GAL population. Negative f_₃_ values indicate potential evidence of admixture, while positive values reflect shared genetic drift. See **Table 1** for population code details.

To better estimate the North African/Middle East component in GAL, we performed a supervised ADMIXTURE analysis using the following reference populations (**Figure 2B**, right): YRI and LWK to represent the sub-Saharan African component, CHB and JPT for Central Asia, and GBR and CEU for Europe. For the North African and Middle East dataset, we selected only those individuals having >90% (in an initial run) or >80% (in a second run) of their primary ancestry component, representing their most genuine North African and Middle East ancestry. This filtering aimed to minimize the impact of their European ancestry on the estimates. Under the 90% threshold, only a few Bedouin genomes (*n* = 18) met the criteria, while a higher number of Bedouin (*n* = 21) and several Mozabite genomes (*n* = 18) passed the 80% filter. In the initial run, GAL shared ancestry proportions of 84.5%, 13.5%, and 2.0% with Europe, Middle East, and sub-Saharan Africa, respectively, compared to 87.0%, 11.6%, and 1.2% in IBS, and 92.2%, 7.8%, and 0.0% in BAS. In the second run, GAL shared ancestry of 83.2%, 7.4%, 8.7% and 0.6% with Europe, Middle East, North Africa and sub-Saharan Africa, respectively, compared to 86.9%, 6.7%, 5.9% and 0.4% in IBS, and 94.9%, 4.0% and 0.0% in BAS. Therefore, the North African/Middle East component in GAL ranged from 13.5% to 16.1% (mean: 14.8%) compared to 7.8% to 12.6% in IBS (mean: 8.25%), and 5.1% and 7.8% in BAS (mean: 6.45%). The differences between ancestry components between these datasets were statistically significant for all comparisons (**Figure S2 – Supplementary File**). In addition, we also detected a minor yet statistically significant Sub-Saharan ancestry to GAL, ranging from 1.0%–2.8% (mean: 1.9%), which is higher than values observed in IBS (0.3%–1.2%; mean: 0.75), and BAS (0%). Interestingly, although the North African and Middle East component is very homogeneous across Galicia, we observed a subtle but clear clinal geographic pattern, with a slightly higher proportion in the southern regions, gradually decreasing toward the North (**Figure 2B**).

According to fastGLOBETROTTER, the estimated admixture contribution from Mozabite (North African-like) ancestry to the GAL dataset is 6.9%, while Middle Eastern-like ancestry accounts for 4.6%, yielding a combined total of 11.5%. This value is slightly lower than the estimates obtained from ADMIXTURE.

To further test the robustness of these admixture inferences, we used the *f_3_*-statistics. As shown in **Figure 2C**, the results of the f_₃_-statistics indicate that the admixture patterns observed in the GAL dataset are statistically compatible with historical gene flow involving various combinations of European and African source populations. Specifically, several *f_3_*(X_EUR_, X_NAF/SSH_; GAL) tests yielded significantly negative values, suggesting that GAL may represent a population formed through admixture between groups genetically related to both European and African ancestries.

Finally, the analysis of mtDNA haplogroups in the GAL dataset reveals that only 1.1% of the samples belong to North African lineages (U6 haplogroups), with no Sub-Saharan lineages (L haplogroups) detected. In sharp contrast, an analysis of the Y-chromosomes of GAL males (*n* = 33) indicates that 21.2% (E1b1b1 haplogroups) show a contribution from North African/Middle Eastern lineages.

### Time Estimates of African Admixture

Time estimates from fastGLOBETROTTER indicate the occurrence of an admixture event in the Galician population that is best explained as gene flow between a predominantly European ancestral group and a more admixed African/Middle Eastern group. The inferred minor source shows a particularly strong North African signature, represented by Mozabites (MOZ), contributing approximately 42%, alongside a smaller Middle Eastern component, represented by Bedouins (BED), accounting for about 19% of the same source. The overall admixture event is dated to approximately 54 generations ago, which translates to a calendar range of around 510–690 CE, assuming a generation time of 26.1 years.

A two-date admixture model was also tested, but it did not provide a better fit than the one-date model, supporting the hypothesis of a single, temporally localized admixture event. The admixture signals were further characterized by the presence of a sub-Saharan African component, with YRI acting as a proxy. While this component was less prominent, it may reflect either a genuine signal of sub-Saharan input or result from model limitations in distinguishing between shared North African and sub-Saharan ancestry.

To validate and complement the findings from fastGLOBETROTTER, admixture dating analyses were also performed using ALDER. In agreement with fastGLOBETROTTER, this software estimated the admixture involving North African ancestry (using MOZ as a proxy) to have occurred approximately 50 generations ago, and admixture with Middle Eastern ancestry (using BED as a proxy) around 54 generations ago. In addition, ALDER identified admixture involving sub-Saharan African ancestry (using YRI and LWK as proxies) at approximately 57 generations ago, which corresponds to around 540 CE.

Taken together, both fastGLOBETROTTER and ALDER yield overlapping time estimates for the introduction of North African and Middle Eastern ancestry into Galicia. For interpretative clarity, we adopt the central point estimates of both methods, 52 generations from ALDER and 54 from fastGLOBETROTTER, to define a reference interval of admixture of 620–670 CE.

We also estimated the timing of the introduction of the North African/Middle Eastern genetic component into various Iberian regions using the NDNAD dataset. While these estimates must be interpreted with caution due to the dataset’s relatively low SNP density and limited sample sizes, they nonetheless reveal informative regional patterns. The inferred admixture dates (in generations, with 95% confidence intervals) were as follows: Andalusia/Central Iberia: 33 (30–35), Catalonia: 39 (34–44), Basque Country/Navarra: 55 (30–70), Cantabria: 46 (40–61), and Asturias: 51 (45–55). Notably, Galicia showed an admixture time of 43 generations in the NDNAD dataset, compared to 52 generations when using the higher-resolution WGS data from the GAL cohort. This discrepancy suggests that the NDNAD estimate may underestimate the true admixture date due to its lower resolution. Despite the inherent limitations, the overall trend across regions supports an earlier admixture in northern Iberia compared to the South and the Center. To further validate this signal, we merged the northern regions (Basque Country/Navarra, Cantabria, Asturias, and Galicia; *n* = 164) to increase statistical power and compared them with the Andalusia/Central Iberia group (*n* = 191). This analysis yielded an admixture estimate of 48 generations (44–51) for northern Iberia, significantly older than the 33 generations estimated for the South-Central regions. These findings suggest that the introduction of North African/Middle Eastern ancestry into Iberia may have occurred in multiple waves or *via* regionally distinct demographic processes, with earlier influxes into the North than previously assumed.

### Fine-Clustering Pattern in Galicia with Reference to National Patterns

At the highest level of genome resolution (WGS), fineSTRUCTURE analysis of the GAL dataset identified 10 terminal branches, though only five formed clusters containing at least four individuals (**Figure 3A**). The Galician genetic landscape is predominantly shaped by a single major cluster, referred to here as the ‘Main’ cluster, comprising 66 genomes and uniformly distributed across most of the region (**Figure 3B**).

**Figure 3.**
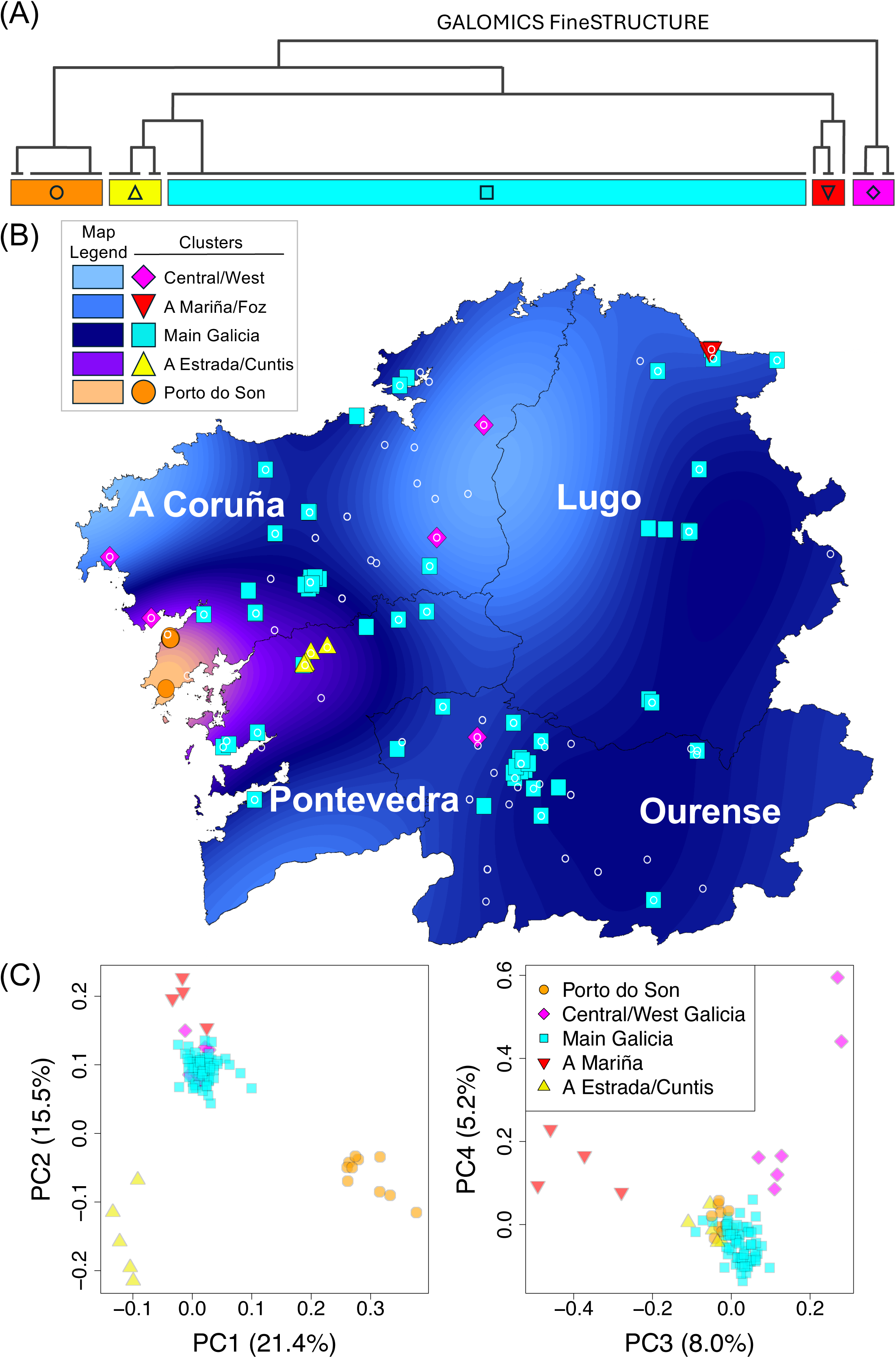
FineSTRUCTURE analysis of GAL whole genomes. (A) Dendrogram depicting the highest level of phylogenetic resolution. (B) PCA of fineSTRUCTURE clusters, illustrating genetic differentiation among individuals. (C) Interpolated frequency map showing the geographic distribution of the identified clusters within the GAL dataset.

The remaining four clusters are significantly smaller and more localized. The ‘Porto do Son’ cluster (West Coast, *n* = 8) shows the most pronounced genetic isolation from the broader Galician population. The ‘Cuntis/A Estrada’ cluster (Inner Pontevedra, *n* = 6) and the ‘A Mariña/Foz’ cluster (northeast Coast, *n* = 4) are also geographically distinct, with the latter located along the northern coastland. Lastly, a small and scattered ‘Central/West’ cluster (*n* = 5) is dispersed throughout central and West Galicia (**Figure 3A and 3B**).

Interpolated frequency maps further emphasize the dominance of the ‘Main’ cluster, while the ‘Porto do Son’ cluster stands out distinctly (**Figure 3B**). The ‘Cuntis/A Estrada’ and ‘A Mariña/Foz’ clusters are less visible due to their blending with the ‘Main’ cluster in their respective areas. Meanwhile, the ‘Central/West’ cluster, being both small and dispersed, is further diluted by the influence of the dominant ‘Main’ cluster. This overall pattern highlights the remarkable homogeneity of the Galician genetic landscape, with only limited instances of localized isolation, most notably in the village of Porto do Son.

The PCA of the fineSTRUCTURE results (**Figure 3C**) also reveals the five clusters described above. PC1 (21.4% variance explained) places the ‘Porto do Son’ and ‘A Estrada//Cuntis’ clusters at the opposite ends of the axis, yet both align on the same pole of PC2 (15.5%), distinguishing them from the remaining clusters. PC3 (8.0%) positions the ‘Central/West Galicia’ and ‘A Mariña’ at opposite extremes, while PC4 (5.2%) further separates these two clusters from the rest. The clustering pattern shown by PC1 to PC4 does not strictly follow the phylogenetic structure suggested by the phylogenetic dendrogram, where, for example, the deepest split occurs between the ‘Central/West’ cluster and the rest. However, it demonstrates a distinct segregation of individuals into five genetically differentiated clusters.

The low genetic structure revealed by WGS data in Galicia contrasts sharply with the highly stratified landscape reported by Bycroft et al. (2019) in their study of the Iberian Peninsula, particularly Galicia, which was based on lower-resolution genomic data. These authors reported identifying 145 genetic clusters among Spanish donors, nearly half of which (∼70) were concentrated in a small, densely populated area within the Galician province of Pontevedra. Remarkably, a closer examination of their data and maps reveals that Galicia accounts for 107 of the 145 clusters (Pardo-Seco *et al*., 2025). Although this specific area of Pontevedra was more densely sampled in Bycroft et al. (2019) than in the GAL study, the observed discrepancy in stratification levels does not appear to be solely attributable to differences in sampling density (Pardo-Seco *et al*., 2025).

In light of the sharp contrast in clustering patterns, we performed a fineSTRUCTURE analysis on the NDNAB dataset (*n* = 453, with 474,761 SNPs after applying the same filters as for the whole-genome data). This dataset includes independent Galician samples alongside donors from various Spanish regions, enabling a detailed assessment of Galicia’s population structure within a national context. This dataset encompasses individuals from Galicia, Asturias, Navarra, Cantabria, the Basque Country, Catalonia, Andalusia, Castilla-La Mancha, and Castilla y León, all regions also examined in Bycroft et al.’s study (2019); **Figure 4A**. Consistent with our findings in the GAL dataset, fineSTRUCTURE identified only three clusters among Galician donors in the NDNAB dataset. Similarly, one cluster contained most of the Galician samples (*n* = 46), mirroring the ‘Main’ GAL cluster, while the remaining two minor clusters comprised 7 and 3 individuals, respectively. In the broader NDNAB Iberian dataset, a total of 23 clusters were detected (including the Galician ones), each closely reflecting the geographic origin of the donors (**Figure 4B**; **Figure S3 – Supplementary File**). Unlike Bycroft et al., (2019) the first major split in the phylogenetic tree of our NDNAB Spanish dataset separates clusters exclusively found in the northern and geographically adjacent region of Cantabria (three clusters) and the Basque Country/Navarra (two clusters) from the rest of the Iberian Peninsula. The next phylogenetic subdivision separates Asturias (six clusters, with one dominant cluster dominating most individuals in the region [*n* = 36]), followed by Catalonia (with a unique and well-differentiated cluster; *n* = 94). Galician genomes emerge at the most recent branching point of the phylogeny, clustering alongside a mixed group of individuals from Andalusia and Central Iberia. This group comprises seven clusters, with three major ones including 128, 35, and 32 individuals, respectively. This clustering pattern differs markedly from that reported by Bycroft et al. (2019), where Galicians from Pontevedra, their hyper-highly stratified Galician province, exhibited the earliest genetic differentiation and accounted for about half of the total clusters found in Iberia. In their analysis, the next major split separated the Basque Country individuals from the rest of the Iberian Peninsula, followed by a division isolating the remaining Galician samples in their dataset.

**Figure 4.**
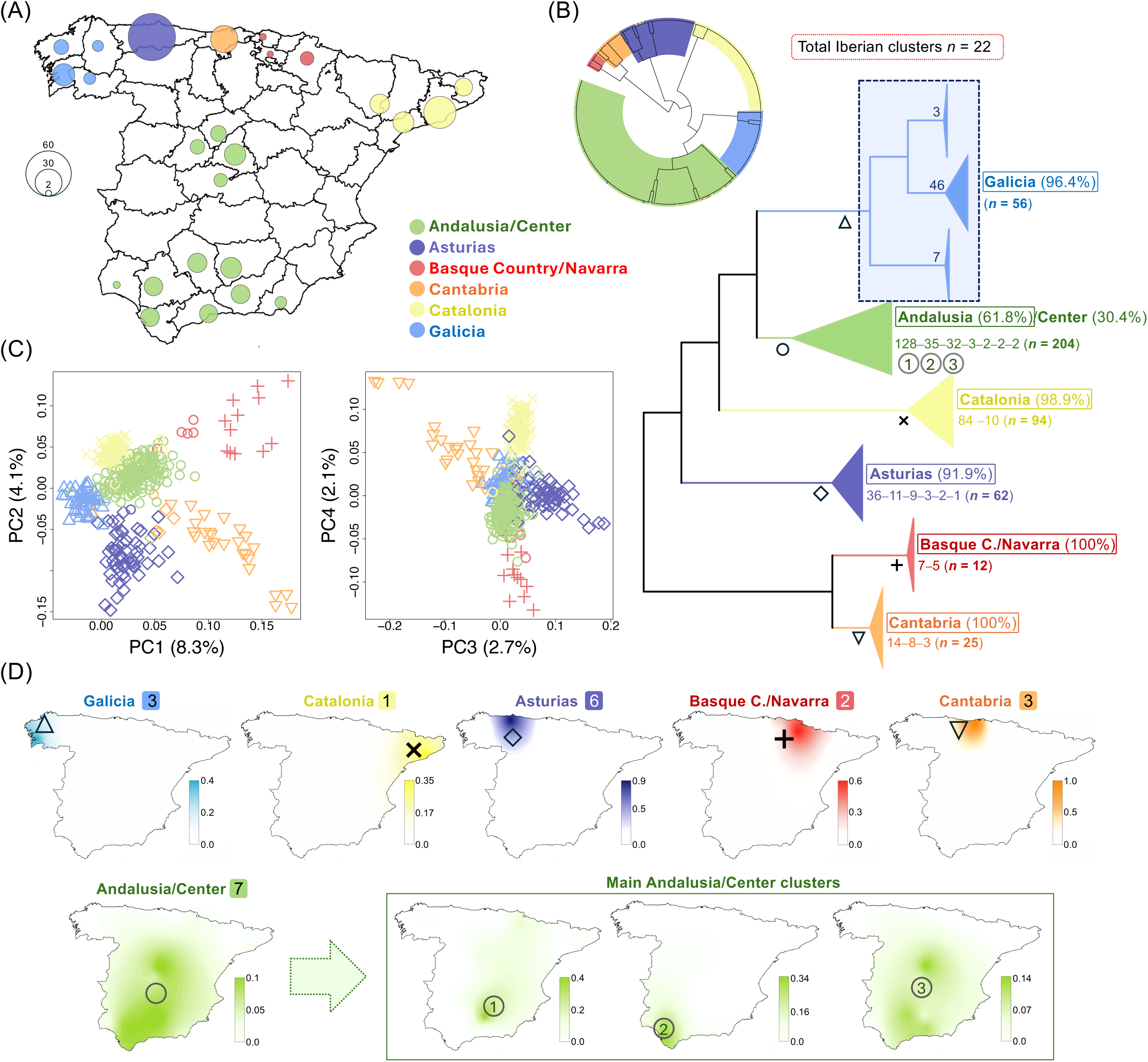
(A) Map illustrating the sampling locations of the NDNAB dataset, which includes Iberian donors whose grandparents were born in the same provinces. Circle sizes are proportional to sample sizes and are centered on the main provincial capital cities. (B) Circular phylogeny (top-left) of the NDNAB dataset (fully expanded in **Figure S3 – Supplementary File**) and a dendrogram (right), where terminal branches are collapsed into nodes representing major geographic regions in Iberia (except Galicia), corresponding to the main autochthonous populations. Unlike other regions, the three Galician clusters were kept separate to facilitate direct comparisons with those observed in the GAL dataset. The proportions indicated to the right of the regional names represent the percentage of samples in each cluster originating from the corresponding autochthonous region, acknowledging that some individuals were sampled in other (often neighboring) regions. Additionally, the total sample size, the number of terminal branches (quantities separated by hyphens), and their respective sample sizes are provided (e.g. fineSTRUCTURE detected three clusters in Cantabria, with sizes 14, 8, and 3 individuals). Symbols in the collapsed nodes of the dendogram correspond to those used in panels C and D. (C) PCA of NDNAB donors, illustrating the clustering pattern observed in (B). (D) Interpolated frequency maps of the continental Spanish territory, depicting the geographic distribution of clusters identified in (B); the numbers following the region names indicate the maximum number of fineSTRUCTURE clusters represented in the maps. For the specific case of the Andalusia/Central region, we show the distribution of the three main clusters in this area (bottom maps), with sample sizes of 128, 35, and 32, each displaying slightly different geographic distributions.

The PCA analysis of the fineSTRUCTURE clusters found in the NDNAB database (**Figure 4C**) further supports the phylogenetic structure observed in the dendrogram (**Figure 4B**). The primary separation in PC1 (8.3%) distinguishes the Cantabria/Basque Country/Navarra clusters from the rest of the dataset. PC2 (4.1%) further differentiates the Basque Country/Navarra from the remaining regions. PC3 (2.7%) highlights the genetic distinction between Asturias and Cantabria, while PC4 (2.1%) again emphasizes the differences among these northern Iberian regions. Notably, all four principal components successfully capture and distinguish the clusters identified in the dendrogram, reinforcing the observed population structure.

### Inbreeding Patterns in GALOMICS

Analyzing inbreeding levels and runs of homozygosity (ROH) segments can reveal events of demographic isolation and offer insights into a population’s demographic history and genetic diversity. This study presents the first investigation of ROH patterns in the Galician population, enabling comparisons with other European and non-European groups at the highest level of genetic resolution possible, namely, that provided by WGS data.

The Galician population (GAL) exhibits ROH patterns comparable to those observed in other European populations (**Figure 5A**). However, a more detailed analysis reveals slightly higher levels of inbreeding compared to the national Spanish dataset represented by the Iberian Peninsula (IBS). Specifically, GAL has a larger count of long ROH segments, longer mean of these segments, and larger cumulative length compared to IBS (*nROH_L_* [GAL] = 9 *vs*. *nROH_L_* [IBS] = 8, *P*-value = 1.5×10^-1^; *mROH_L_* [GAL] = 2.4 *vs*. *mROH_L_* [IBS] = 2.1, *P*-value = 5.2×10^-8^; *sROH_L_*[GAL] = 20.6 *vs*. *sROH_L_* [IBS] = 15.2, *P*-value = 9.2×10^-4^); **Table S2**. In line with these findings, GAL also has a lower count of short ROH segments, a shorter mean of these segments, and a smaller cumulative length compared to IBS (*nROH_S_* [GAL] = 2504 *vs*. *nROH_S_* [IBS] = and 2592, *P*-value = 5.8×10^-19^; *mROH_S_*[GAL] = 0.199 *vs*. *mROH_S_* [IBS] = 0.203, *P*-value = 9.2×10^-19^; *sROH_S_* [GAL] = 499 *vs*. *sROH_S_* [IBS] = 525, *P*-value = 3.4×10^-26^); **Table S2**. Overall, these metrics support the observations of significantly higher inbreeding levels in GAL relative to IBS and other European datasets, except when comparing long ROH segments in historically isolated populations (Basques and Sardinians), which show higher values. Additionally, total inbreeding coefficient values indicate higher inbreeding in GAL compared to IBS (*F_ROH_* [GAL] = 0.007 *vs*. *F_ROH_* [IBS] = 0.005, *P*-value = 9.3×10^-4^) and other continental European datasets, although still lower than in Sardinians and Basques; **Table S2**.

**Figure 5.**
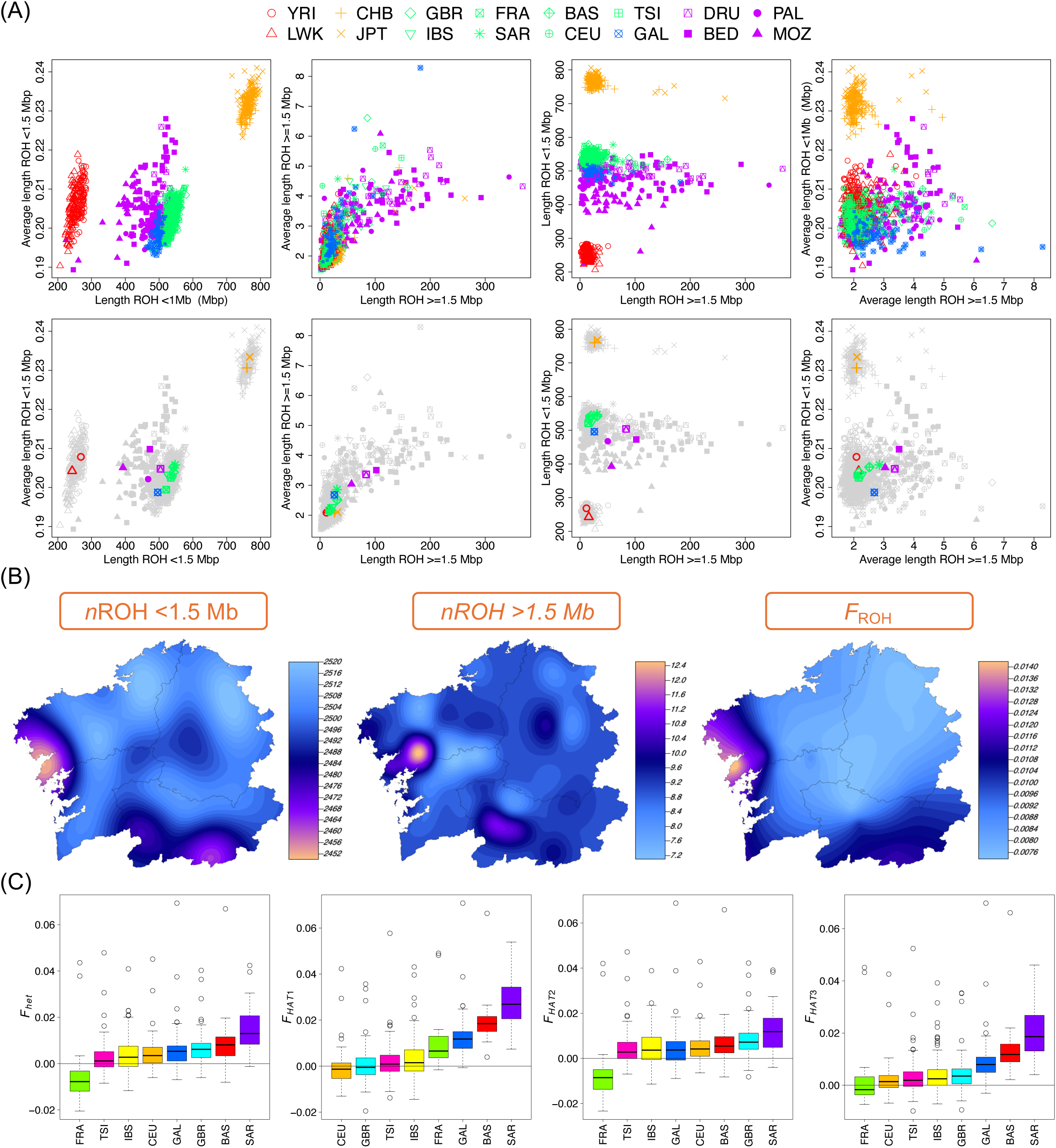
Analysis of inbreeding in GAL genomes and reference worldwide population datasets. (A) Various ROH estimates indicate that GAL shows patterns consistent with those expected for a dataset of predominantly European ancestry. (B) Interpolated frequency maps showing the distribution of different ROH values across GAL genomes. (C) F-statistic analysis conducted on European ancestry datasets. Further details on the different statistical measures are provided in **Table S1** and **S2.**

In agreement with ROH values, Galicia (GAL) has higher values of *F_HET_, F_HAT1_* and, *F_HAT3_* compared to IBS (*F_HET_*[GAL] *=* 0.005 *vs. F_HET_* [IBS] = 0.003*, P*-value = 6.4×10^-2^; ; *F_HAT1_* [GAL] = 0.012 *vs. F_HAT1_* [IBS] =.0001*, P*-value = 2.7×10^-16^; and *F_HAT3_* [GAL] = 0.008 *vs. F_HAT3_* [IBS] =.0002*, P*-value = 5.2×10^-9^); and equal values of *F_HAT2_* for both population (*F_HAT2_*[GAL and IBS] = 0.004); **Figure 5C**, **Table S3**. Within Europe, GAL’s metrics are intermediate, with Sardinians and Basques showing the highest homozygosity values among these datasets.

Additionally, the ‘Porto do Son’ cluster significantly contributes to the higher inbreeding values in GAL compared to IBS. For instance, metrics for long ROH segments in this cluster are substantially higher (and statistically significant) than those for the general GAL: *nROH_L_* [Porto do Son] = 11; *mROH_L_* [Porto do Son] = 3.5; and *sROH_L_* [Porto do Son] = 45). Conversely, short ROH segment metrics are generally lower: *nROH_S_* [Porto do Son] = 2442; *mROH_S_*[Porto do Son] = 0.199; *sROH_S_* [Porto do Son] = 487; **Table S2**. In fact, the inbreeding values for the GAL dataset excluding ‘Porto do Son’, are similar to those observed in the other clusters (GAL: *nROH_L_* [GAL without Porto do Son] = 9; *mROH_L_*[GAL without Porto do Son] = 2.6; and *sROH_L_* [GAL without Porto do Son] = 26; *nROH_S_*[GAL without Porto do Son] = 2502; *mROH_S_* [GAL without Porto do Son] = 0.199; *sROH_S_* [GAL without Porto do Son] = 497).

Overall, these results indicate that inbreeding levels in Galicia are broadly comparable to those observed elsewhere on the Iberian Peninsula, with only marginally elevated values in certain localities, such as Porto do Son.

### Regional Variations of Potential Biomedical Interest

To assess allele differences of potential biomedical relevance, we performed a single-site association test comparing the GAL cohort to the IBS dataset. This analysis was restricted to SNPs with a minor allele frequency (MAF) > 0.05 in cases and controls, as the focus was solely on common variants. A total of 85 SNP variants (out of >17M) surpassed the Bonferroni threshold (**Figure S4 – Supplementary File**; **Table S1**). Among these, only one SNP (rs9917044) was located in an exonic region (*ZNF28*; Zinc Finger Protein 28 gene) and was synonymous. Additionally, 25 variants were found in intronic regions, 13 were non-coding but classified as regulatory, and 46 were intergenic. When these variants were compared against a merged European cohort (CEU, GBR, IBS, and TSI) and using a MAF > 0.05 in cases and controls, 57 remained significant after applying the Bonferroni threshold. None of the 85 SNP variants appears as associated with a disease trait in SNPedia. The gene-based test between GAL and IBS identified only four genes as statistically significant after Bonferroni correction: *ATP5F1AP10* (two SNPs), *SDR42E1P4* (two SNPs), *FRG2EP* (17 SNPs), and *DUX4L34* (13 SNPs); only the first two genes (supported on only two SNPs) remained significant when compared to the merged European cohort (**Figure S5 – Supplementary File**).

### Microgeographic Stratification of Disease Risk

By computing Polygenic Risk Scores (PRS) for eleven common diseases, we investigated whether demographic particularities in Galicia, compared to the broader Iberian (IBS) population, influence the distribution of alleles associated with disease risk in GAL genomes. For most PRS metrics, no statistically significant differences were observed between GAL and IBS. However, despite their small quantitative effect, the differences were statistically significant for PRS_ASD_ (IBS median = −0.22 [IQR: −0.68–0.53], GAL median = 0.15 [IQR: −0.41–0.71]; Wilcoxon test *P*-value = 0.038) and PRS_IBD_ (IBS median = −0.25 [IQR: −0.89–0.50], GAL median = 0.25 [IQR: −0.62–0.86]; *P*-value = 0.007); **Figure S6 – Supplementary File**. The wide IQR in these metrics suggests substantial variation among individuals, indicating that some donors have markedly higher or lower polygenic risk relative to others in the group.

To further investigate the microgeographic stratification of disease risk within Galicia, we analyzed the spatial distribution patterns of PRS metrics for the eleven common diseases examined across the region. Rather than being homogeneously distributed, the maps in **Figure 6A** illustrate distinct geographic patterns of disease risk. Notably, ovarian cancer risk is more pronounced along the western Atlantic coastline, peaking in the southwestern coastal region. In contrast, breast cancer risk (although slightly varying depending on the PRS used) is generally elevated in both eastern and western Galicia but lower in a central North-to-East longitudinal band. Coronary artery disease and inflammatory bowel disease exhibit broadly similar geographic distributions. Atrial fibrillation risk follows a pronounced South-to-North gradient, with lower risk in the South and higher risk in the North. Type 2 diabetes risk peaks in central Galicia and reaches its lowest values in the southeastern corner. Acute lower respiratory infection risk is lowest in the northwestern corner and highest in a small northeastern region. Autism spectrum disorder exhibits intermediate risk values from mid-central to northern Galicia, with the lowest risk in the southwest and the highest in the southeast. Schizophrenia risk is highest in the southeasternmost corner of Galicia and remains elevated across most of the region, except for the northwestern corner. In contrast, Alzheimer’s disease follows an approximately inverse pattern.

**Figure 6.**
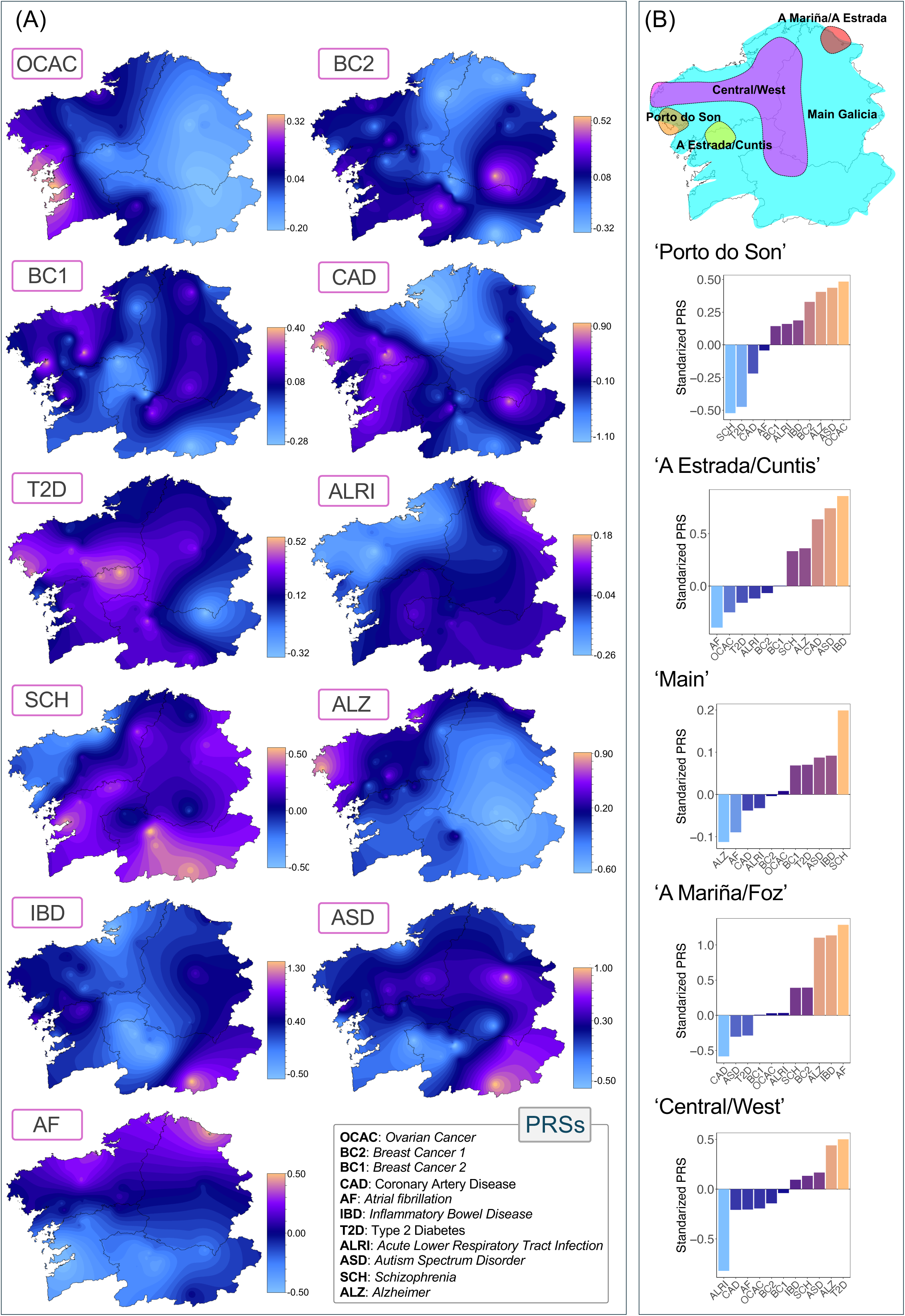
Interpolated maps showing the distribution of polygenic risk for common diseases across the Galician territory. (B) Median PRS values for various common diseases across the fineSTRUCTURE clusters, ordered from lowest to highest risk. Note that median values do not fully capture the variability of PRS distributions (which can be seen in detail in **Figure S7 – Supplementary File**) and may not directly correspond to the interpolated maps in (A). These estimates are preliminary and may vary with larger sample sizes in future analyses, but they establish a foundation for further academic exploration. The top panel in (B) provides a graphical representation of the geographic locations of the main Galician clusters, as inferred from Figure 3.

Motivated by these findings, we further examined disease risk stratification by genetic cluster. Although the sample sizes for most clusters (except for the ‘Main’ cluster) were limited, and the different PRS scores encompass a wide range of variant counts (ranging from dozens to millions), we identified statistically significant differences in several comparisons (**Figure S7 – Supplementary File**). The most notable findings include: (a) Coronary artery disease risk: Significant differences were observed between ‘Porto do Son’ and ‘A Estrada/Cuntis’ (*P*-value = 0.016), ‘A Estrada/Cuntis’ and ‘A Mariña/Foz’ (*P*-value = 0.019), and ‘A Estrada/Cuntis’ and the rest of Galicia (*P*-value = 0.038); (b) Inflammatory bowel disease risk: Significant differences were found between ‘A Estrada/Cuntis’ and the ‘Main’ cluster (*P*-value = 0.026), as well as between ‘A Estrada/Cuntis’ and the rest of Galicia (*P*-value = 0.040). Additionally, differences were observed between the ‘Main’ cluster and ‘A Mariña/Foz’ (*P*-value = 0.025), and between ‘A Mariña/Foz’ and the rest of Galicia (*P*-value = 0.038); (c) Atrial fibrillation risk: Significant differences were detected between the ‘Main’ cluster and ‘A Mariña/Foz’ (*P*-value = 0.007), and between ‘A Mariña/Foz’ and the rest of Galicia (*P*-value = 0.008); (d) Alzheimer’s disease risk: Differences were observed between the ‘Main’ cluster and ‘A Mariña/Foz’ (*P*-value = 0.012), the ‘Main’ cluster and the rest of Galicia (*P*-value = 0.007), and between ‘A Mariña/Foz’ and the rest of Galicia (*P*-value = 0.027); and (e) Autism spectrum disorder risk: A significant difference was found between ‘A Estrada/Cuntis’ and ‘A Mariña/Foz’ (*P*-value = 0.019). Therefore, of the 13 statistically significant contrasts identified, 8 involved the ‘A Mariña/Foz’ cluster and 6 involved the ‘A Estrada/Cuntis’ cluster. Despite their relatively small sample sizes, these two clusters account for a substantial proportion of the major disease risk stratification observed in our Galician sample. In addition, PRS metrics for the different clusters can be ranked according to the diseases considered. For instance, the ‘Porto do Son’ cluster has the highest risk for ovarian cancer and the lowest for schizophrenia, the ‘A Mariña/Foz’ cluster has the highest risk for atrial fibrillation but the lowest for coronary artery disease, while the ‘A Estrada/Cuntis’ cluster has the lowest risk for atrial fibrillation and the highest risk for inflammatory bowel disease (**Figure 6B**).

## Discussion

Galicia’s geographic position at the far western edge of the Iberian Peninsula, combined with its historical isolation, has led some studies to hypothesize that the region may harbor unique genomic features (Brage *et al*., 2007; Bycroft *et al*., 2019; Esperón-Moldes *et al*., 2019; Fachal *et al*., 2012; Fernández-Marmiesse *et al*., 2006; Salas *et al*., 1998). For the first time, whole-genome sequencing (WGS) data from Galicia allow an in-depth investigation at the highest possible genomic resolution, revealing patterns of genetic variation that are broadly shared with other Iberian populations, yet also display some distinct regional traits. The genetic landscape of the Galician population (GAL) reflects a demographic history consistent with early settlement, subtle differentiation from the broader Iberian Peninsula, and an early admixture with North African and Middle Eastern ancestries. Principal component analysis (PCA) and fineSTRUCTURE clustering reveal that Galicians form specific but closely related genetic clusters, showing only limited divergence from other Iberian groups.

### Lack of Evidence for Long-Term Genetic Isolation

Galicia demonstrates slightly higher values of *nROH_L_*, *sROH_L_*, and *mROH_L_*, compared to IBS and other European datasets, with statistically significant differences observed after Bonferroni correction. This pattern extends to shorter ROH segments, resulting in a higher total inbreeding coefficient (*F_ROH_*) than in IBS and other Europeans. In agreement with these findings, inbreeding coefficients indicate slightly higher overall homozygosity in Galicia compared to IBS and other European datasets. While statistically significant differences with the rest of the Peninsula exist, they are quantitatively modest and largely driven by the distinct “Porto do Son” cluster (as also inferred from PCA plots and spatial genetic maps).

The slightly higher inbreeding coefficients in Porto do Son likely reflect historical geographic isolation and elevated consanguinity. Marriage patterns within small coastal or rural communities, combined with limited gene flow, can increase genetic homogeneity over generations. Nevertheless, even in Porto do Son, inbreeding levels remain within the Iberian norm, indicating that Galicia’s demographic and cultural factors produce autozygosity patterns similar to those of neighboring regions. Therefore, the slightly elevated inbreeding levels observed in Galicia challenge the hypothesis of pronounced population stratification in the region (against Bycroft et al. (2019)). If strong stratification were present, one would expect allele frequency disparities within Galicia to result in numerous genetic clusters and elevated homozygosity levels consistent with a Wahlund effect, a pattern not observed in this study. The minor divergence existing between GAL and the rest of the Iberian Peninsula most likely resulted from genetic drift and small-scale fluctuations in allele frequencies across the genome. There is no compelling evidence to suggest that long-standing geographic isolation is the primary factor shaping the genetic landscape of Galicia.

The relatively elevated African ancestry component existing in Galicia (see discussion below), greater than in other parts of Iberia, may have played a role in mitigating levels of inbreeding through increased outbreeding, thereby contributing to the preservation of genetic diversity in the Galician population.

### Minimal Population Structure

The limited number of clusters inferred by fineSTRUCTURE from Galician whole genomes (ten in total, or which only five included more than four individuals) and the moderate levels of inbreeding observed in the GAL population suggest a demographic history marked by extensive gene flow, both within Galicia and from genetically distinct, non-Galician populations, including groups from North African and Middle East. Such a pattern aligns with Galicia’s geographic and socio-historical context, characterized by: (*i*) the absence of major geographical barriers and (*ii*) a predominantly rural, dispersed demographic structure. These factors likely promoted both intra-regional mobility and external genetic exchange, helping to preserve high genetic diversity and limiting inbreeding across the relatively small geographical area of Galicia. While certain areas in Galicia may have experienced localized demographic conditions conducive to increased inbreeding or mild founder effects, as previously documented for mutations causing autosomal recessive congenital ichthyosis (Fachal *et al*., 2012) and other inherited disorders, our data reveal a markedly lower degree of fine-scale population structure than that reported by Bycroft et al. (2019). Their study, based on low-density genome-wide data, identified ∼70 genetic clusters recognized by these authors. However, as critically reanalyzed in Pardo-Seco et al. (2025), the actual number of clusters in their data appears to exceed 107 across the Galician territory out of 145 observed in the whole Iberian Peninsula, accounting for ∼74% of the total. These clusters are heavily concentrated in a small southwestern section of Pontevedra province, near the border with northern Portugal. This strikingly high number of clusters is particularly unexpected given the province’s relatively small size, its high population density (Pontevedra is among the most densely populated provinces in Spain, with over 200 inhabitants per square kilometer; https://ide.depo.gal), and its historical, cultural, and religious significance. Notably, the city of Tui in this area has long served as a key religious center in both Galicia and northern Portugal. The discrepancy is even more pronounced considering that this southwestern region is one of the most cosmopolitan areas in Galicia, home to Vigo (the province’s largest urban center and Galicia’s most populous city), and one of the world’s busiest fishing ports. Moreover, the region lacks any substantial geographical barriers that might explain deep population substructure. As critically discussed by Pardo-Seco et al. (2025), the sharp discrepancies between our results and those in Bycroft et al. (2019) could have been due to technical artifacts, amplified by the high sensitivity of the fine-STRUCTURE tool, as well as flawed misinterpretations of the data.

### The Riddle of African Ancestry in Galicia

Several studies (most of them focusing on uniparental markers) suggest that Galicia has a proportion of North African genetic ancestry that is comparable to or even exceeds that of other Iberian regions (**Supplementary Text**), despite its geographic distance from North Africa. Using high-resolution WGS data, we estimate that the North African/Middle East genomic component in GAL ranges from 13.5% to 16.1%, significantly higher than previously reported for autosomal data and exceeding estimates for other Iberian datasets (IBS: 7.8%–12.6%; BAS: 5.1%–7.8%). Approximately half of this component is shared with “Mozabite-like” ancestry, with Mozabites (MOZ) representing Berbers from Algeria’s M’Zab Valley. The remaining proportion is attributed to Middle Eastern ancestry, primarily represented by the Bedouin (BED) population, nomadic Arab groups historically associated with the Syrian and Arabian deserts, who expanded throughout the Levant, Mesopotamia, and North Africa following the Islamic conquests of the 7th century. This African ancestry is present throughout the entire Galician territory, implying long-term, extensive gene flow and demographic integration across the region. When examining the spatial distribution of this component using interpolated individual frequencies, a subtle yet consistent South-to-North gradient of North African and Middle Eastern ancestry emerges, suggesting a southern entry point into Galicia *via* maritime or overland routes from the southwest. This pattern warrants further investigation in future studies with larger sample sizes and comparable spatial and genomic resolution. A novel and noteworthy observation is that the North African component is much more pronounced in the Y-chromosome (21.2%) and autosomal genome (13.5%–16.5%) than in the mitochondrial genome (1.0%), suggesting a sex-biased admixture event with a predominant paternal contribution. This male-skewed pattern is consistent with broader trends observed throughout the Iberian Peninsula, likely reflecting the historical context of migrations before, during, or after the Islamic expansion, where male-dominated incursions and settlements were common.

Moreover, the detection of sub-Saharan ancestry may suggest an even more complex demographic dynamic than the North African/Middle Eastern– European admixture process. This could include either actual (direct) sub-Saharan input or model misattribution due to shared drift or admixture history between ‘Mozabites-like’ ancestry and sub-Saharan groups like YRI. In our analysis, we detected a low but statistically significant sub-Saharan African component in both GAL (1.0%–2.8%; mean: 1.9%) and IBS (0.3%–1.2%; mean: 0.75), whereas this component was absent in the BAS dataset. Given that the reference populations used in this study are essentially unadmixed, this sub-Saharan component remains undetectable when analyzed within North African or sub-Saharan datasets. Furthermore, the minimal representation of this component in Iberian datasets, combined with the limited availability of comprehensive African WGS data, presents a challenge for further investigation into its precise origins. The existence of this sub-Saharan component in Iberia has been particularly reported in the literature on uniparental markers. A meta-analysis of mtDNA variation in Iberian by Barral-Arca et al. (2016) identified the presence of a characteristic sub-Saharan component in various Iberian regions, with proportions ranging from 0% to 5% (e.g., Asturias: 3.6%; Galicia: 2.9%; La Rioja: 5.0%; with a mean of 1.4% in mainland Spain). The introduction of typical sub-Saharan genetic variation into Europe, particularly in Iberia, has been explored by Cerezo et al. (2012). Phylogeographic analyses of mtDNA profiles suggest that approximately 65% of European L-lineages (typically indicative of sub-Saharan mtDNA African ancestry) likely arrived in Iberia during historical periods, including Romanization, Arab rule, and the Atlantic slave trade. The remaining 35% of European-specific subclades of L mtDNA may have originated from gene flow between sub-Saharan Africa and Europe as early as 11,000 years ago. Additionally, the presence of sub-Saharan mtDNA components has been documented in Iberian Romani populations (Gómez-Carballa *et al*., 2013). More recently, ancient DNA studies further support the introgression of early sub-Saharan populations into Iberia, providing evidence of an early prehistoric migration route from Africa to the Iberian Peninsula, likely *via* the Strait of Gibraltar (González-Fortes *et al*., 2019).

The admixture signals detected by GLOBETROTTER and ALDER point to a historically significant episode of gene flow into Galicia, likely occurring between 620 and 670 CE. The predominant African-related ancestry in this event appears to derive from North African populations, with a smaller but noteworthy Middle Eastern component. The presence of sub-Saharan genetic signatures, though more modest, may reflect earlier contact or ancestral gene flow mediated through North African intermediary populations, consistent with patterns of trans-Saharan connectivity. Moreover, our admixture timing estimates reveal a striking pattern within the Iberian Peninsula: the North African and Middle Eastern genetic components appear to have entered northern Iberia significantly earlier than in central and southern regions. Notably, after accounting for the potential underestimation of admixture dates (as inferred from comparisons between high- and low-resolution genomic data in Galicia), the pooled northern regions (including the Basque Country/Navarra, Cantabria, Asturias, and Galicia) show an average admixture date around 770 CE (range: 690–880 CE). In contrast, estimates for Andalusia and central Iberia are more recent, averaging around 1160 CE (range: 1110–1240 CE).

While the southern admixture signal might align with the well-documented Arab-Berber expansion during the Islamic conquest and subsequent centuries of rule, the earlier dates in the North (though requiring cautious interpretation) challenge this conventional narrative and suggest more complex or independent episodes of contact and gene flow. Overall, these findings highlight a notable North African/Middle Eastern genetic influence on the Iberian Peninsula, particularly in the North, and more specifically in the northwestern region of Galicia, that appears to predate the advent of Islamic rule. While our data estimate the arrival of this component in a single, concise admixture event occurring 620 and 670 CE, it is possible that statistical limitations, methodological constraints, or, more likely, the demographic complexity of this ancestral signal obscure multiple layers of gene flow that may have begun as early as the Roman period or even in (pre-)historic times (Cerezo *et al*., 2012; González-Fortes *et al*., 2019). One could, for instance, hypothesize a steady, small-scale migration, trade networks, and/or multifaceted influx of this ancestry into Galicia over several centuries and predating the Islamic conquest, with a peak occurring during the 6th to 7th century, giving rise to the signal detected in our model. Investigating the connections between North Africa/Middle East and Galicia during the pre-Islamic period is essential to understanding this genetic legacy, especially considering the documented limited influence of the Islamic era in northern Iberia.

Thus, numerous historical links between North Africa and Galicia can already be traced back to the Roman era. One such link is evident in the case of soldiers from the *Cohors I Celtiberorum*, who were initially stationed in the Roman province of *Mauretania Tingitana* (in present-day northern Morocco) in 132 CE (at the end of Trajan’s reign), and were later posted to *Cidadela*, in Sobrado dos Monxes (A Coruña, Galicia), where a garrison of ∼500 members was accommodated. Likely, many of these soldiers settled locally upon discharge (Caamaño-Gesto, 2015; Costa-García, 2009), and the cohort remained stationed in *Cidadela* until around the 4th century. Through this demographic movement, it is plausible that some of its members may have introduced African ancestry to Galicia. More importantly, in Roman times, military units like the *Cohors I Celtiberorum* rarely moved in isolation: they were typically accompanied by a civilian population comprising families, merchants, craftsmen, and service providers. This civilian presence supported the logistical needs of the soldiers and facilitated the establishment of semi-permanent settlements near military camps. There is evidence of these civilian population settlements, referred to as *vicus*, *insulae,* or even *vicus*, in *Cidadela*. Curiously, a nearby town still preserves the name ’Insua,’ possibly reflecting this historic and demographic legacy. It is therefore plausible that accompanying North African civilian groups contributed to the local demographic and genetic landscape, facilitating not only military but also cultural, demographic, and economic exchanges between North Africa and northwestern Iberia.

Additionally, there is evidence of well-established Atlantic trade networks, centered on salted fish, garum, and especially cereals transported in amphorae (e.g. from present-day Tunez), that connected North African ports to the Galician coast *via* the Balearic Islands, the Iberian Levant, and the Strait of Gibraltar, bringing with them merchants, sailors, and laborers of North African origin. (Fernández Fernández *et al*., 2021; Fernández-Fernández, 2014; Naveiro Lopez, 1991). Although much of this commerce may have had a primarily cultural rather than demographic impact, these intertwined military and commercial currents might help to establish a North African genetic legacy in Galicia centuries before any Islamic political or cultural influence.

Another example from Roman times is the presence of the *Legio VII Gemina*, established in the North of Iberia (part of the *Gallaecia* province), which included an important presence of people originating in Africa (Palao-Vicent, 1998). In addition, there is abundant historical documentation for the Late Antique period concerning the North African and Middle Eastern influence in Galicia, covering the span from the end of Roman rule in the northwest of Iberia to the onset of the Islamic era (i.e., 5th to 7th centuries CE). During this time, the Iberian Peninsula underwent significant transformations following the collapse of Roman authority and the emergence of new political and religious structures, particularly under the Visigothic Kingdom. While slavery had long been a feature of Roman society, there is considerable textual evidence that the slave trade not only persisted but intensified during this period, often taking on new religious and economic dimensions. Churches, Monasteries, and civil elites, especially in northwestern Iberia (modern-day Galicia), became prominent economic and spiritual centers after the 5th century (Díaz-Martínez, 1987). These institutions, especially monasteries and churches, relied heavily on enslaved labor for agriculture, construction, and domestic service. Ecclesiastical institutions and the aristocracy were large landowners who needed manpower, most of whom were slaves, designated as *servi* in the texts of the time. Around 655 CE, *Richimirus*, Bishop of Dume, who preceded *Fructuosus* as head of the monastery, bequeathed in his testament the liberation and emancipation of numerous *servi*, and to these he donated >500 slaves, some of whom came from the property of the deceased bishop (Díaz-Martínez, 1987; Maciel, 1996). This source provides insight into the massive number of slaves that may have been arriving in the northwest during the 6th and especially the 7th centuries. By the 6th and 7th centuries, the Iberian Peninsula was linked to broader Mediterranean/Atlantic trade and slave routes, including those connecting North Africa and the Eastern Mediterranean. Slaves of African or Middle Eastern origin were often brought into Iberia *via* maritime and overland routes (through Carthage, Ceuta, or southern Hispania). This route operated in both directions, as indicated by historical texts: many slaves were Britons and Saxons who were sold in markets in Rome and Constantinople (Fernández-Fernández, 2014). These enslaved individuals may have been prisoners of war, victims of raids, or traded through expanding networks controlled by Byzantine, Berber, or Visigothic intermediaries. Contemporary texts once again inform us about slaves who were brought to Gaul to supply the needs of churches and other major landowners, including individuals of Mauritanian origin (Pieri, 2005). Evidence suggests that specialized traders, possibly of Eastern origin, such as Syrians, Greeks, and Jews, dominated this market, organizing systematic raids to capture slaves in the Mauritanian territories, beyond the reach of contemporary political and administrative control (Pieri, 2005). It is also plausible that a significant community of Eastern merchants settled in the area during the 6th and 7th centuries. This community, likely connected to the Visigothic Church and monarchy, oversaw the flow of goods between the Mediterranean and Galicia and may have played a key role in managing this highly profitable slave trade (Fernández-Fernández, 2014; Tereso *et al*., 2015).

Finally, the recent genomic analysis of the remains of the alleged Bishop Teodomiro of Iria Flavia (Galicia), born toward the end of the 7^th^ century, indicates the presence of an important North African component (∼15%, Pérez-Ramallo, personal communication). The authors of the study state that “…their ancestors may have received North African gene flow during the Roman period or, more recently, during or after the Islamic conquest” (Pérez-Ramallo *et al*., 2024); see also (Tereso *et al*., 2015). The recent findings from Oteo-García et al. (2025) also align with the need to conceive a significant pre-Islamic genetic impact on the Iberian Peninsula. These authors reconstructed the medieval genetic history of eastern Iberia using ancient DNA from the Valencian region. Their analysis, which focused on the effects of the Morisco deportations, shows that North African– related ancestry was already present in medieval Valencia and remained largely unchanged following the Reconquista.

Taken together, this body of evidence challenges the traditional view that North African and Middle Eastern genetic influence in northern Iberia resulted solely from a South-to-North diffusion during Islamic rule and the *Reconquista*; *contra* (Bycroft *et al*., 2019). Instead, it suggests that much earlier, independent episodes of gene flow may have contributed to this genetic legacy. Potential factors shaping the demographic landscape of northwestern Iberia include pre-historic contacts, movements during the Roman period, post-Phoenician trade networks, and trans-Mediterranean contacts predating the 8th-century conquest, including the trafficking of enslaved individuals. Religious institutions, particularly monasteries, likely played a central role in this process, acting both as recipients and administrators of enslaved labor.

### Patterns of variation in Galicia or biomedical interest

Despite the limited statistical power, no specific common genetic markers of biomedical significance were identified that distinctly differentiate Galicia from the broader Iberian Peninsula. A gene-based analysis using SKAT, which accounts for rare variation, identified a small number of genes with statistically significant differences compared to the IBS cohort. Additionally, minor differences in disease risk between GAL and IBS were observed for various common diseases; however, the magnitude of these differences remains relatively low.

Of more interesting consideration is our observation of the disease risk micro-geographically stratified within Galicia. We demonstrate that disease risk follows distinct spatial patterns, with certain regions exhibiting elevated risk for specific diseases while showing lower risk for others. Furthermore, for the first time, we illustrate how disease risk stratifies by genetic cluster, an approach that does not necessarily align with the patterns observed in PRS interpolated geographical maps. Notably, we show that it is possible to rank disease risk within genetic clusters, offering a novel perspective on the distribution of common disease susceptibility across the region.

While our observation of regional differences in PRS values for various conditions in Galicia is academically relevant, its implications for public health policy remain uncertain. On the one hand, PRS reflects genetic risk alone and constitutes just one of many factors contributing to overall disease susceptibility. For instance, in breast and ovarian cancer, risk stratification algorithms incorporate additional variables such as hormone replacement therapy, parity, endometriosis, contraceptive use, and age of menarche. At the regional level, where Galicia has full autonomy in managing public health policies, the primary focus would likely be on optimizing individual risk stratification through appropriate PRSs, rather than on accounting for microgeographic variation within the region. Incorporating such fine-scale differences would be particularly challenging for public health authorities due to factors such as population mobility and the complexity of tracing ancestral origins. For example, while our study includes individuals with all four grandparents born within an 80 km radius, real-world clinical cases are likely to exhibit more diverse regional ancestries. Moreover, the prevailing trend in the scientific community favors the development of PRSs with broad applicability across large populations, as seen with the widely used CanRisk model for breast and ovarian cancer. This trend is likely to hold even in populations with significantly more complex ancestry patterns than Galicia. For example, efforts are underway to develop software capable of improving PRS portability by integrating fine-scale ancestry components into PRS predictive power, as exemplified by ANCHOR, which has been applied to UK Biobank data (Hu *et al*., 2025).

Moreover, although subtle differences in SNP frequencies and gene markers are observed between Galicians and the broader Iberian population, the magnitude of these differences appears limited, casting doubt on the cost-effectiveness of implementing large-scale, region-specific WGS initiatives that are not explicitly focused on disease-related applications. If allele frequencies, linkage disequilibrium patterns, and genetic risk predictions remain largely consistent between Galicia and the broader Iberian region, and if existing national and international PRS models already demonstrate strong performance in predicting disease risk among Galicians, the rationale for extensive regional WGS projects becomes questionable. Instead, considering the still substantial costs associated with WGS, and even more critically, the challenges related to the storage and management of such voluminous data, a more targeted genomic approach, such as focusing sequencing efforts on coding regions through custom gene panels or WES, may offer a more efficient and cost-effective alternative. This strategy would be particularly valuable for prioritizing conditions of high public health relevance, such as hereditary breast and colorectal cancers. Supporting this view, recent research from Regeneron Pharmaceuticals (Gaynor et al., 2024) based on UK Biobank data indicates that prioritizing imputed WES over full WGS can substantially improve the discovery rate in genetic association studies, delivering critical insights while optimizing resource allocation.

## Conclusions

This study represents the first comprehensive analysis of WGS variation in the Galician population. Despite Galicia’s historical reputation as a genetically isolated region within the Iberian Peninsula, our results reveal no strong signatures of long-term isolation. On the contrary, we observed low levels of fine-scale population structure, challenging previous claims of extreme substructure in the area.

Importantly, we identified a significantly higher North African and Middle Eastern genetic component in Galician genomes than previously reported. This ancestry is distributed across the entire Galician territory, suggesting widespread historical admixture and sustained gene flow within the region. Such connectivity may have reduced levels of inbreeding that might otherwise be expected in a small, demographically scattered population, particularly when contrasted with other Iberian groups with larger effective population sizes. We detected a male-biased contribution to this African component, along with a subtle South-to-North frequency gradient. This geographic pattern may reflect a southern entry point into Galicia, possibly *via* maritime or overland routes, though further investigation is needed to confirm this hypothesis. Most notably, the timing of admixture predates Islamic rule in the Iberian Peninsula, challenging the traditional assumption that the observed African component in Galicia is primarily the result of Arab-Berber expansion and the subsequent *Reconquista*.

In addition, our analyses show that PRS vary modestly across Galicia, with evidence of microgeographic stratification both regionally and among fine-scale genetic clusters. However, we also found that WGS provides limited additional resolution over exome sequencing for the characterization of common, clinically relevant variants in this population. This highlights the importance of matching sequencing strategies to specific research goals. In many cases, targeted approaches such as exome or gene panel sequencing may offer a more cost-effective means of maximizing clinically actionable insights.

## Author’s contributions

ASE and F.M-T conceived the project and funding. JPS, ACM, XB, AGC, LCM, CMC, and ASE carried out the analyses. ACM and XB designed the GALOMICS web resource. JPS, AGC, CMC, and ASE contributed to a critical discussion of the main findings. ASE wrote the draft of the study. All the authors critically review the final version of the manuscript.

## Acknowledgements

We would like to thank all the members of the GALOMICS Worlding Group collaborators for their invaluable help in sampling recruitment. We also kindly like to thank historians and archaeologists David Peterson, Olalla López Costas, Patxi Pérez Ramallo, Ramón Fábregas Valcarce, Roberto Pena Puentes, Rubén Álvarez Asorey, and particularly, Adolfo Fernández Fernández, for their valuable insights during discussions on the historical aspects of this study.

## Supplementary Data

**Supplementary File**. Complementary figures to those included in the main text.

**Table S1**. Top candidate variants exceeding the Bonferroni threshold in the single-point association test comparing GAL vs. IBS, along with corresponding values from the same test comparing GAL vs. a merged European dataset.

**Table S2**. ROH statistics computed on worldwide datasets.

**Table S3.** F-statistic analysis conducted on European ancestry datasets.

## GALOMICS Working Group – GenPoB & GenViP IDIS research groups

Alba Camino Mera, Alberto Gómez Carballa, Alicia Carballal Fernández, Ana Cotovad Bellas, Ana Isabel Dacosta Urbieta, Ana María Pastoriza Mourelle, Ana María Senín Ferreiro, Andrés Efraín Muy Pérez, Antía Rivas Oural, Antonio Justicia, Grande, Antonio Piñeiro García, Antonio Salas Ellacuriaga, Anxela Cristina Delgado García, Ángela Manzanares Casteleiro, Belén Mosquera Pérez, Blanca Díaz Esteban, Carlos Durán Suárez, Carmen Curros Novo, Carmen Rodríguez-Tenreiro Sánchez, Conrado Martínez Cadenas, Cristina Serén Trasorras, Cristina Talavero González, Esther Montero Campos, Federico Martinón-Torres, Fernando Álvez González, Fernando Caamaño Viña, Gloria Viz Rodríguez, Hugo Alberto Tovar Velasco, Irene Rivero Calle, Irene Visos Varela, Isabel Ferreirós Vidal, Isabel Rego Lijo, Iván Prieto Gómez, Jacobo Pardo Seco, Jesús Eirís Puñal, José Gómez Rial, José Manuel Fernández García, José María Martinón Sánchez, Julia García Currás, Julián Montoto Louzao, Lara Martínez Martínez, Laura Navarro Marrón, Lidia Piñeiro Rodríguez, Lorenzo Redondo Collazo, Lúa Castelo Martínez, Manuel Vázquez Donsión, María Elena Gamborino Caramés, María Elena Sobrino Fernández, María José Currás Tuala, María Martínez Castro, María Soledad Vilas Iglesias, María Sol Rodriguez Calvo, María Sonia Ares Gómez, María Teresa Autrán García, Marina Casas Pérez, Marta Aldonza Torres, Marta Bouzón Alejandro, Marta Lendoiro Fuentes, Miguel Sadiki Orayyou, Miriam Ben García, Miriam Cebey López, Miriam Taboada Puga, Monserrat López Franco, Narmeen Mallah, Natalia García Sánchez, Nour El Zahraa Mallah, Ouhao Zhu Huang, Regueiro Casuso, Rosaura Picáns Leis, Ruth Barral Arca, Sandra Carnota Antonio, Sandra Viz Lasheras, Sara Pischedda, Sara Rey Vázquez, Silvia Cerqueiro Rey, Sonia Marcos Alonso, Sonia Serén Fernández, Susana María Rey García, Ulises Toscanini, Vanesa Álvarez Iglesias, Wiktor Dominik Nowak, Xabier Bello Paderne

## References

Alexander, D. H., Novembre, J. & Lange, K. (2009). Fast model-based estimation of ancestry in unrelated individuals. Genome Res 19, 1655–64.

Amigo, J., Salas, A. & Phillips, C. (2011). ENGINES: exploring single nucleotide variation in entire human genomes. BMC Bioinformatics 12, 105.

Amigo, J., Salas, A., Phillips, C. & Carracedo, Á. (2008). SPSmart: adapting population based SNP genotype databases for fast and comprehensive web access. BMC Bioinformatics 9, 428.

Barral-Arca, R., Pischedda, S., Gómez-Carballa, A., Pastoriza, A., Mosquera-Miguel, A., López-Soto, M., et al. (2016). Meta-Analysis of Mitochondrial DNA Variation in the Iberian Peninsula. PLoS One 11, e0159735.

Brage, A., Tomé, S., Garcia, A., Carracedo, Á. & Salas, A. (2007). Clinical and molecular characterization of Wilson disease in Spanish patients. Hepatol Res 37, 18–26.

Butler-Laporte, G. & Povysil, G. & Kosmicki, J. A. & Cirulli, E. T. & Drivas, T. & Furini, S., et al. (2022). Exome-wide association study to identify rare variants influencing COVID-19 outcomes: Results from the Host Genetics Initiative. PLoS Genet 18, e1010367.

Bycroft, C., Fernandez-Rozadilla, C., Ruiz-Ponte, C., Quintela, I., Carracedo, A., Donnelly, P., et al. (2019). Patterns of genetic differentiation and the footprints of historical migrations in the Iberian Peninsula. Nat Commun 10, 551.

Byrne, R. P., Martiniano, R., Cassidy, L. M., Carrigan, M., Hellenthal, G., Hardiman, O., et al. (2018). Insular Celtic population structure and genomic footprints of migration. PLoS Genet 14, e1007152.

Caamaño-Gesto, J. M. (2015). Marcas de la Cohors I Celtiberoum halladas enel campamento romado de Cidadela (Sobrado dos Monxes, A Coruña). Portvgalia, Nova Série 36, 107–120.

Camino-Mera, A., Pardo-Seco, J., Bello, X., Argiz, L., Boyle, R. J., Custovic, A., et al. (2024). Whole Exome Sequencing Identifies Epithelial and Immune Dysfunction-Related Biomarkers in Food Protein-Induced Enterocolitis Syndrome. Clin Exp Allergy.

Cann, H. M., Toma, C. d., Cazes, L., Legrand, M.-F., Morel, V., Piouffre, L., et al. (2002). A Human Genome Diversity Cell Line Panel. Science 296, 261–262.

Cavalli-Sforza, L. L. (2005). The Human Genome Diversity Project: past, present and future. Nat Rev Genet 6, 333–40.

Cerezo, M., Achilli, A., Olivieri, A., Perego, U. A., Gómez-Carballa, A., Brisighelli, F., et al. (2012). Reconstructing ancient mitochondrial DNA links between Africa and Europe. Genome Res 22, 821–826.

Chang, C. C., Chow, C. C., Tellier, L. C., Vattikuti, S., Purcell, S. M. & Lee, J. J. (2015). Second-generation PLINK: rising to the challenge of larger and richer datasets. Gigascience 4, 7.

Conrad, O., Bechtel, B., Bock, M., Dietrich, H., Fischer, E., Gerlitz, L., et al. (2015). System for automated geoscientific analyses (SAGA) v. 2.1. 4. Geoscientific Model Development 8, 1991–2007.

Costa-García, J. M. (2009). Behind Cohors I Celtiberorum: the archaeological evidence. Universidad de Valladolid, Spain.

Díaz-Martínez, P. d. l. C. (1987). Formas económicas y sociales en el monacato visigodo. Ediciones Universidad de Salamanca, Salamanca.

Dopazo, J., Amadoz, A., Bleda, M., García-Alonso, L., Alemán, A., García-García, F., et al. (2016). 267 Spanish Exomes Reveal Population-Specific Differences in Disease-Related Genetic Variation. Mol Biol Evol 33, 1205–18.

Esperón-Moldes, U. S., Pardo-Seco, J., Montalván-Suárez, M., Fachal, L., Ginarte, M., Rodríguez-Pazos, L., et al. (2019). Biogeographical origin and timing of the founder ichthyosis TGM1 c.1187G > A mutation in an isolated Ecuadorian population. Sci. Rep. 9, 7175.

Ewels, P., Magnusson, M., Lundin, S. &Kaller, M. (2016). MultiQC: summarize analysis results for multiple tools and samples in a single report. Bioinformatics 32, 3047–8.

Fachal, L., Rodriguez-Pazos, L., Ginarte, M., Toribio, J., Salas, A. &Vega, A. (2012). Multiple local and recent founder effects of *TGM1* in Spanish families. PLoS One 7, e33580.

Fernández Fernández, A., Rodríguez Nóvoa, A. A., Valle Abad, P. &Ruanova Álvarez, N. (2021). Roman fish salting factory in praia do Naso (Illa de Arousa, Pontevedra). Minius 26, 137–161.

Fernández-Fernández, A. (2014). El comercio tardoantiguo (ss. IV-VI) en el noroeste peninsular a través del registro cerámico de la Ría de Vigo. Archaeopress, Oxford.

Fernández-Marmiesse, A., Salas, A., Vega, A., Fernández-Lorenzo, J. R., Barreiro, J. &Carracedo, Á. (2006). Mutation spectra of *ABCC8* gene in Spanish patients with Hyperinsulinism of Infancy (HI). Hum Mutat 27, 214.

Fiorito, G., Di Gaetano, C., Guarrera, S., Rosa, F., Feldman, M. W., Piazza, A., et al. (2016). The Italian genome reflects the history of Europe and the Mediterranean basin. Eur J Hum Genet 24, 1056–62.

Garcia-Calleja, J., Biagini, S. A., de Cid, R., Calafell, F. & Bosch, E. (2025). Inferring past demography and genetic adaptation in Spain using the GCAT cohort. Sci Rep 15, 14225.

Genome of the Netherlands, C. (2014). Whole-genome sequence variation, population structure and demographic history of the Dutch population. Nat Genet 46, 818–25.

Genomes Project, C., Auton, A., Brooks, L. D., Durbin, R. M., Garrison, E. P., Kang, H. M., et al. (2015). A global reference for human genetic variation. Nature 526, 68–74.

Gilbert, E., O’Reilly, S., Merrigan, M., McGettigan, D., Vitart, V., Joshi, P. K., et al. (2019). The genetic landscape of Scotland and the Isles. Proc Natl Acad Sci U S A 116, 19064–19070.

Gómez-Carballa, A., Pardo-Seco, J., Fachal, L., Vega, A., Cebey, M., Martinón-Torres, N., et al. (2013). Indian signatures in the westernmost edge of the European Romani diaspora: new insight from mitogenomes. PLoS One 8, e75397.

González-Fortes, G., Tassi, F., Trucchi, E., Henneberger, K., Paijmans, J. L. A., Díez-del-Molino, D., et al. (2019). A Western route of prehistoric human migration from Africa into the Iberian Peninsula. *Proc. Royal Soc. A* 286, 20182288.

Grove, J., Ripke, S., Als, T. D., Mattheisen, M., Walters, R. K., Won, H., et al. (2019). Identification of common genetic risk variants for autism spectrum disorder. Nat Genet 51, 431–444.

Hartl, D. L.Clark, A. G. (2007). Principles of populatin genetics (4th ed.). Sinauer Associates, Sunderland.

Hu, S., Ferreira, L. A. F., Shi, S., Hellenthal, G., Marchini, J., Lawson, D. J., et al. (2025). Fine-scale population structure and widespread conservation of genetic effect sizes between human groups across traits. Nat Genet 57, 379–389.

Karakachoff, M., Duforet-Frebourg, N., Simonet, F., Le Scouarnec, S., Pellen, N., Lecointe, S., et al. (2015). Fine-scale human genetic structure in Western France. Eur J Hum Genet 23, 831–6.

Kerminen, S., Havulinna, A. S., Hellenthal, G., Martin, A. R., Sarin, A. P., Perola, M., et al. (2017). Fine-Scale Genetic Structure in Finland. G3 (Bethesda) 7, 3459–3468.

Lawson, D. J., Hellenthal, G., Myers, S. & Falush, D. (2012). Inference of population structure using dense haplotype data. PLoS Genet 8, e1002453.

Lee, A., Mavaddat, N., Wilcox, A. N., Cunningham, A. P., Carver, T., Hartley, S., et al. (2019). BOADICEA: a comprehensive breast cancer risk prediction model incorporating genetic and nongenetic risk factors. Genet Med 21, 1708–1718.

Lee, S., Emond, M. J., Bamshad, M. J., Barnes, K. C., Rieder, M. J., Nickerson, D. A., et al. (2012). Optimal unified approach for rare-variant association testing with application to small-sample case-control whole-exome sequencing studies. Am J Hum Genet 91, 224–37.

Lee, S.Miropolsky, L. (2023). SKAT: SNP-Set (Sequence) Kernel Association Test - R package version 2.2.5.

Leslie, S., Winney, B., Hellenthal, G., Davison, D., Boumertit, A., Day, T., et al. (2015). The fine-scale genetic structure of the British population. Nature 519, 309–314.

Li, H. (2013). Aligning sequence reads, clone sequences and assembly contigs with BWA-MEM. *arXiv:1303.3997v2 [q-bio.GN]*.

Li, H.Durbin, R. (2010). Fast and accurate long-read alignment with Burrows-Wheeler transform. Bioinformatics 26, 589–95.

Loh, P. R., Lipson, M., Patterson, N., Moorjani, P., Pickrell, J. K., Reich, D., et al. (2013). Inferring admixture histories of human populations using linkage disequilibrium. Genetics 193, 1233–54.

Maciel, M. J. (1996)*. Libro antiguidade tardia e paleocristianismo em Portugal*. Pedidos, Edic ões Colibri, Lisboa.

Mathieson, I.McVean, G. (2012). Differential confounding of rare and common variants in spatially structured populations. Nat Genet 44, 243–6.

Monti, R., Eick, L., Hudjashov, G., Lall, K., Kanoni, S., Wolford, B. N., et al. (2024). Evaluation of polygenic scoring methods in five biobanks shows larger variation between biobanks than methods and finds benefits of ensemble learning. Am J Hum Genet 111, 1431–1447.

Moorjani, P., Patterson, N., Hirschhorn, J. N., Keinan, A., Hao, L., Atzmon, G., et al. (2011). The history of African gene flow into Southern Europeans, Levantines, and Jews. PLoS Genet 7, e1001373.

Naveiro Lopez, J. L. (1991)*. El comercio antiguo en el N.W. Peninsular*. Museo arqueolóxico, A Coruña.

Oteo-Garcia, G., Silva, M., Foody, M. G. B., Yau, B., Fichera, A., Alapont, L., et al. (2025). Medieval genomes from eastern Iberia illuminate the role of Morisco mass deportations in dismantling a long-standing genetic bridge with North Africa. Genome Biol 26, 108.

Palao-Vicent, J. J. (1998). La participación de africani en la Legio VII Gemina. *Iberia* 1.

Pardo-Seco, J., Gómez-Carballa, A., Martinez-Cadenas, C., Martinón-Torres, F. & Salas, A. (2025). Challenging patterns of fine-scale genetic structure in Galicia and East-to-West genetic divergence across the Iberian Peninsula: When technical batches rewrite history. *in preparation*.

Pardo-Seco, J., Llull, C., Berardi, G., Gómez, A., Andreatta, F., Martinón-Torres, F., et al. (2016). Genomic continuity of Argentinean Mennonites. Sci. Rep. 6, 36392.

Pardo-Seco, J., Viz-Lasheras, S., Bello, X., Gomez-Carballa, A., Camino-Mera, A., Pischedda, S., et al. (2024). Whole exome sequencing identifies new susceptibility candidates underlying community-acquired pneumonia. Genes Dis 11, 101170.

Patterson, N., Price, A. L. &Reich, D. (2006). Population structure and eigenanalysis. PLoS Genet 2, e190.

Pérez-Ramallo, P., Rodríguez-Varela, R., Staniewska, A., Ilgner, J., Krzewińska, M., Chivall, D., et al. (2024). Unveiling Bishop Teodomiro of Iria Flavia? An attempt to identify the discoverer of St James’s tomb through osteological and biomolecular analyses (Santiago de Compostela, Galicia, Spain). Antiquity 98, 973–990.

Pieri, D. (2005)*. Le commerce du vin oriental à l’Époque byzantine (V-VII siècles)*. Institut Français du Proche-Orient, Beirut.

Poplin, R., Chang, P. C., Alexander, D., Schwartz, S., Colthurst, T., Ku, A., et al. (2018). A universal SNP and small-indel variant caller using deep neural networks. Nat Biotechnol 36, 983–987.

Ralf, A., Gonzalez, D. M., Zhong, K. & Kayser, M. (2018). Yleaf: Software for Human Y-Chromosomal Haplogroup Inference from Next-Generation Sequencing Data. Mol Biol Evol 35, 1820.

Rentzsch, P., Schubach, M., Shendure, J. & Kircher, M. (2021). CADD-Splice-improving genome-wide variant effect prediction using deep learning-derived splice scores. Genome Med 13, 31.

Salas, A., Comas, D., Lareu, M. V., Bertranpetit, J. & Carracedo, Á. (1998). mtDNA analysis of the Galician population: a genetic edge of European variation. Eur J. Hum. Genet. 6, 365–75.

Salas, A., Pardo-Seco, J., Barral-Arca, R., Cebey-López, M., Gómez-Carballa, A., Rivero-Calle, I., et al. (2018). Whole Exome Sequencing identifies new host genomic susceptibility factors in empyema caused by *Streptococcus pneumoniae* in children: A pilot study. Genes (Basel) 9.

Salas, A., Pardo-Seco, J., Cebey-López, M., Gómez-Carballa, A., Obando-Pacheco, P., Rivero-Calle, I., et al. (2017). Whole Exome Sequencing reveals new candidate genes in host genomic susceptibility to Respiratory Syncytial Virus Disease. Sci. Rep. 7, 15888.

Serra-Vidal, G., Lucas-Sanchez, M., Fadhlaoui-Zid, K., Bekada, A., Zalloua, P. & Comas, D. (2019). Heterogeneity in Palaeolithic Population Continuity and Neolithic Expansion in North Africa. Curr Biol 29, 3953–3959 e4.

Tereso, S., Brito, A. F., Umbelino, C., Cipriano, M., André, C. & Carvalho, P. C. (2015)*. Arquelogía funerária alto medieval da Torre Velha (Castro de Avelãs, Gragança)*. Bilbao : Universidad del País Vasco / Euskal Herriko Unibertsitatea.

The R Core Team (2019). R: A Language and Enviroment for Statistical Computing. R Foundation for Statistical Computing, Vienna, Austria.

Valls-Margarit, J., Galvan-Femenia, I., Matias-Sanchez, D., Blay, N., Puiggros, M., Carreras, A., et al. (2022). GCAT|Panel, a comprehensive structural variant haplotype map of the Iberian population from high-coverage whole-genome sequencing. Nucleic Acids Res 50, 2464–2479.

Wang, R. J., Al-Saffar, S. I., Rogers, J. & Hahn, M. W. (2023). Human generation times across the past 250,000 years. Sci Adv 9, eabm7047.

Wangkumhang, P., Greenfield, M. & Hellenthal, G. (2022). An efficient method to identify, date, and describe admixture events using haplotype information. Genome Res 32, 1553–1564.

Weissensteiner, H., Pacher, D., Kloss-Brandstätter, A., Forer, L., Specht, G., Bandelt, H.-J., et al. (2016). HaploGrep 2: mitochondrial haplogroup classification in the era of high-throughput sequencing. Nucleic Acids Res. 44, W58–W63.

Yun, T., Li, H., Chang, P. C., Lin, M. F., Carroll, A. & McLean, C. Y. (2021). Accurate, scalable cohort variant calls using DeepVariant and GLnexus. Bioinformatics 36, 5582–5589.

Zheutlin, A. B., Dennis, J., Karlsson Linner, R., Moscati, A., Restrepo, N., Straub, P., et al. (2019). Penetrance and Pleiotropy of Polygenic Risk Scores for Schizophrenia in 106,160 Patients Across Four Health Care Systems. Am J Psychiatry 176, 846–855.

