## Supplementary File for "Whole-genome sequencing in Galicia reveals male-biased pre-Islamic North African ancestry, subtle population structure, and micro-geographic patterns of disease risk"

Complementary figures to those included in the main text.

**Figure S1**. Number of inhabitants living in the donor's and their grandparents' localities of origin, according to official data (<https://www.ige.gal/>).

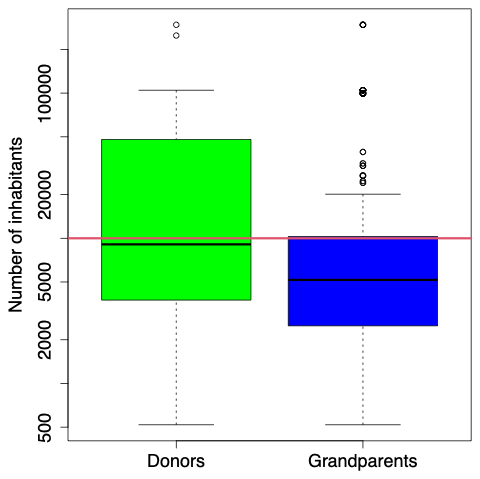

**Figure S2**. Percentages of ancestry in Galicia according to the different models of ADMIXTURE: *K* = 4, unsupervised; *K* = 4 unsupervised ; K = 5 supervised; and *K* = 5 supervised and 90% filter of ancestry for the main ancestral component in North Africa and Near East.

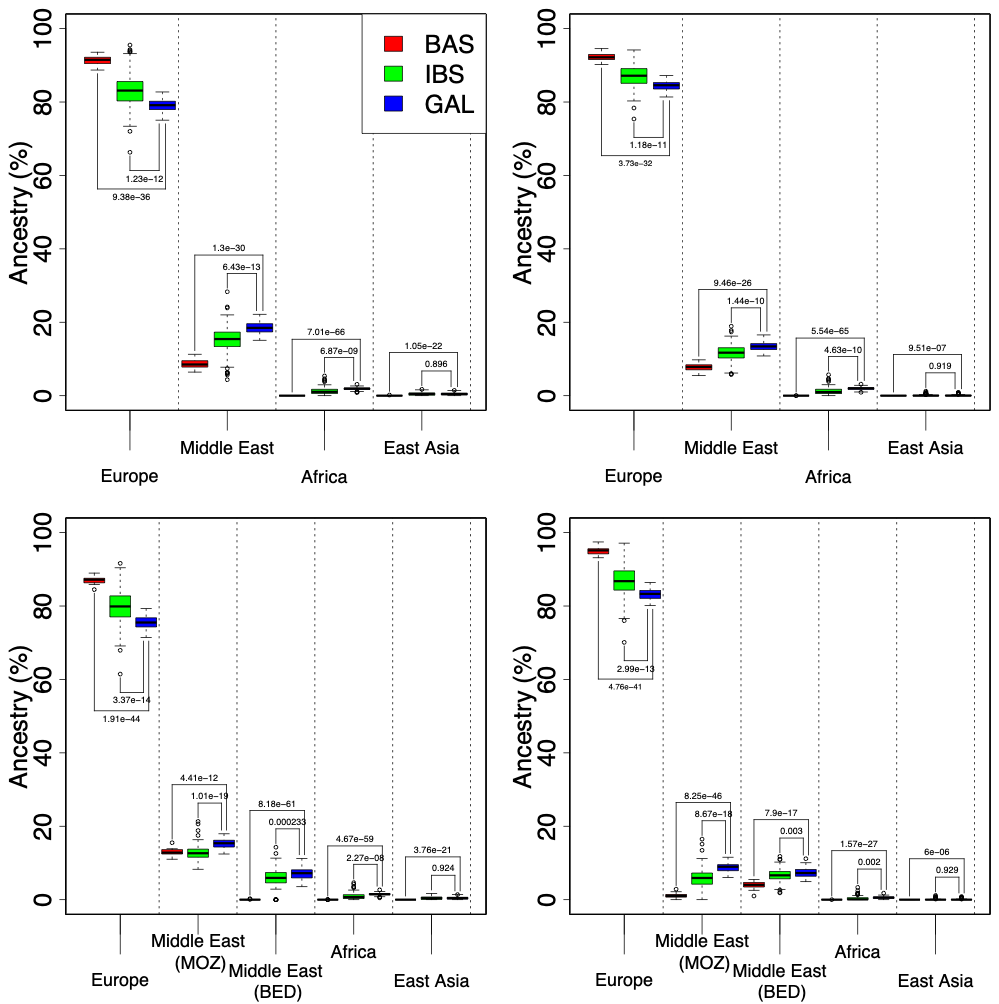

**Figure S3.** Phylogeny of fineSTRUCTURE clusters inferred from the Iberian NB database. This geographically detailed phylogeny (at the provincial level) corresponds to the scheme presented in **Figure 4B** of the main document.

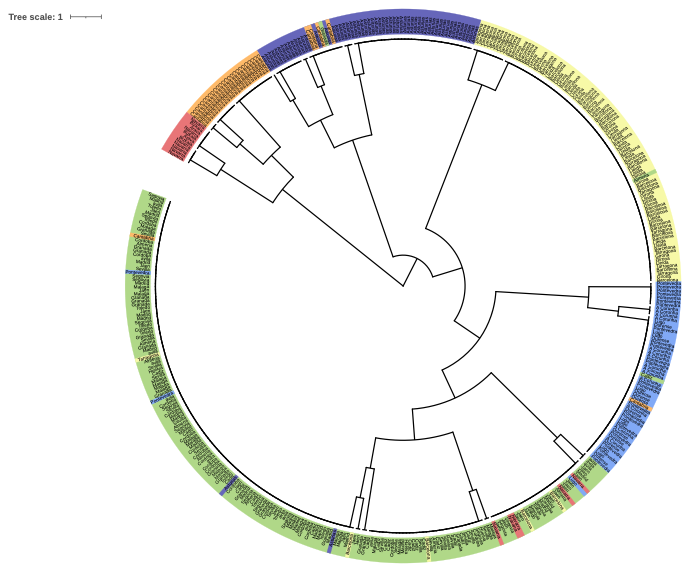

**Figure S4**. Mahatam plot (A) and QQ-plot (B) of the single-point association test between SNPs in GAL and IBS.

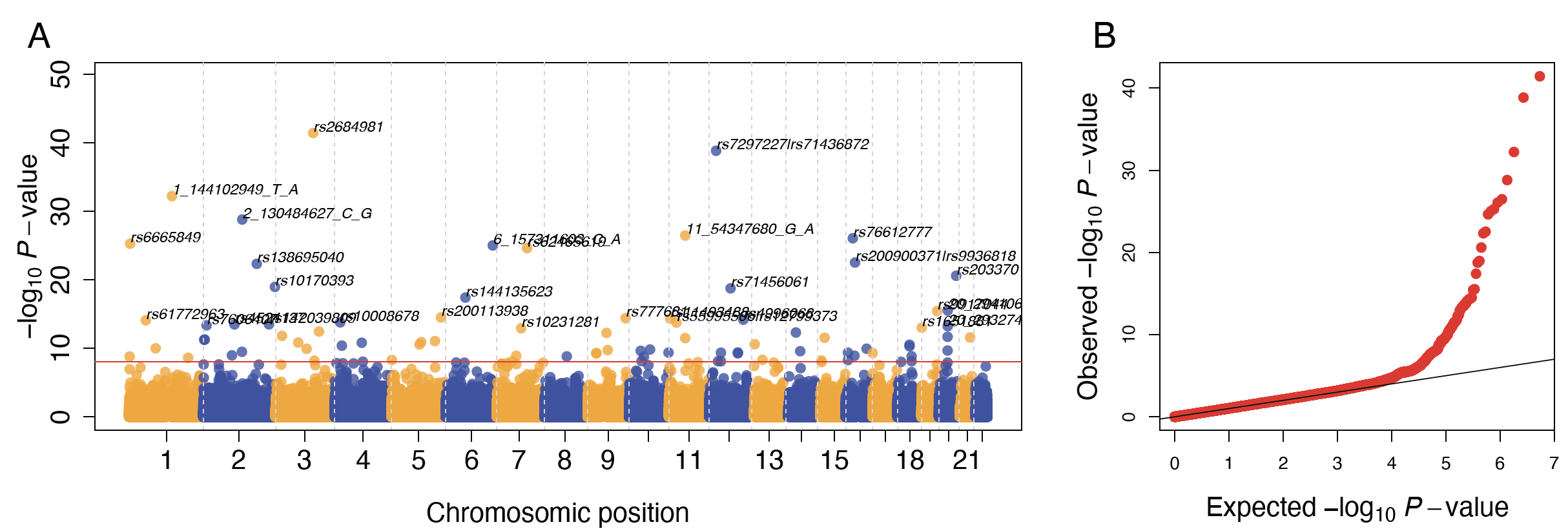

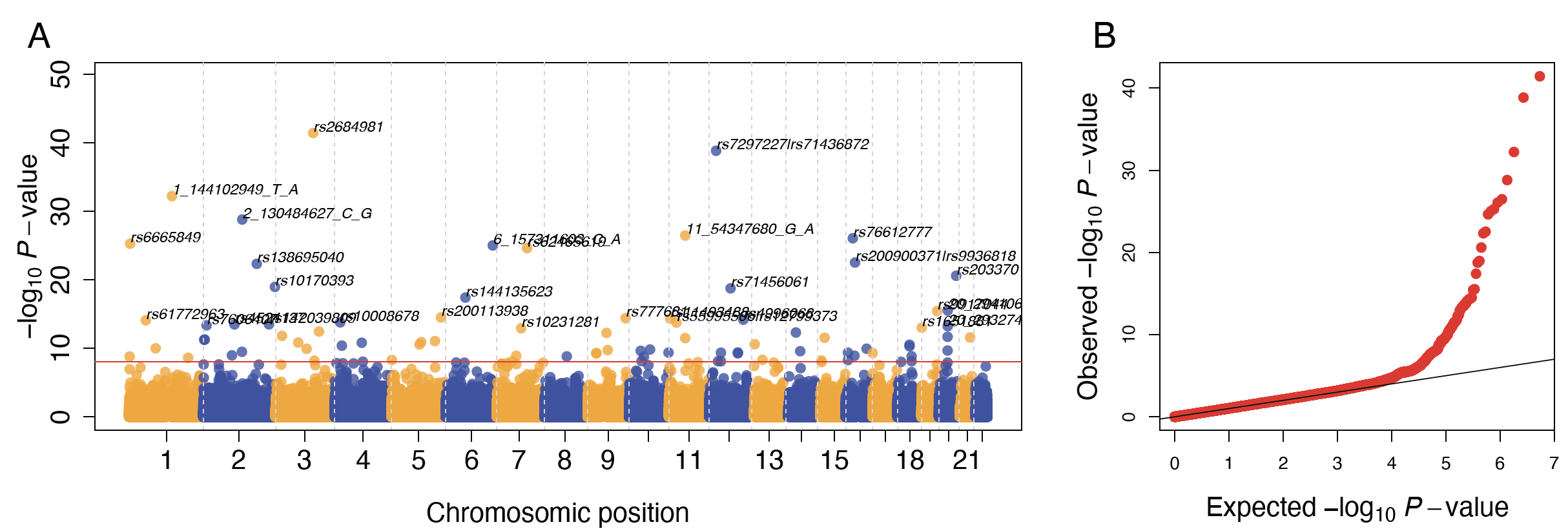

**Figure S5**. Mahatam plot (A) and QQ-plot (B) of gene-based SKAT-O association test between SNPs in GAL and IBS.

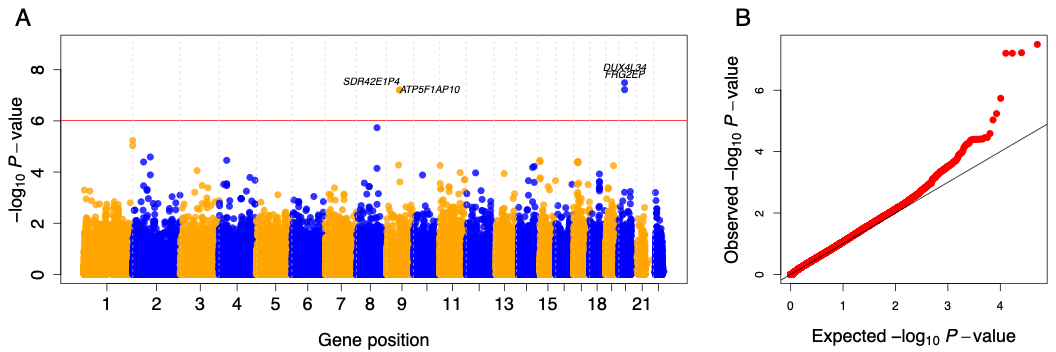

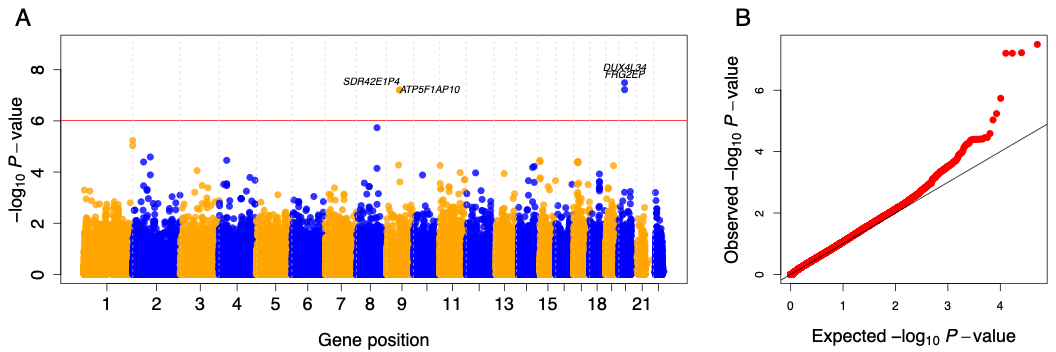

**Figure S6**. Boxplots of PRS values for different common diseases in GAL and IBS population datasets. On the right, median values and IQR.

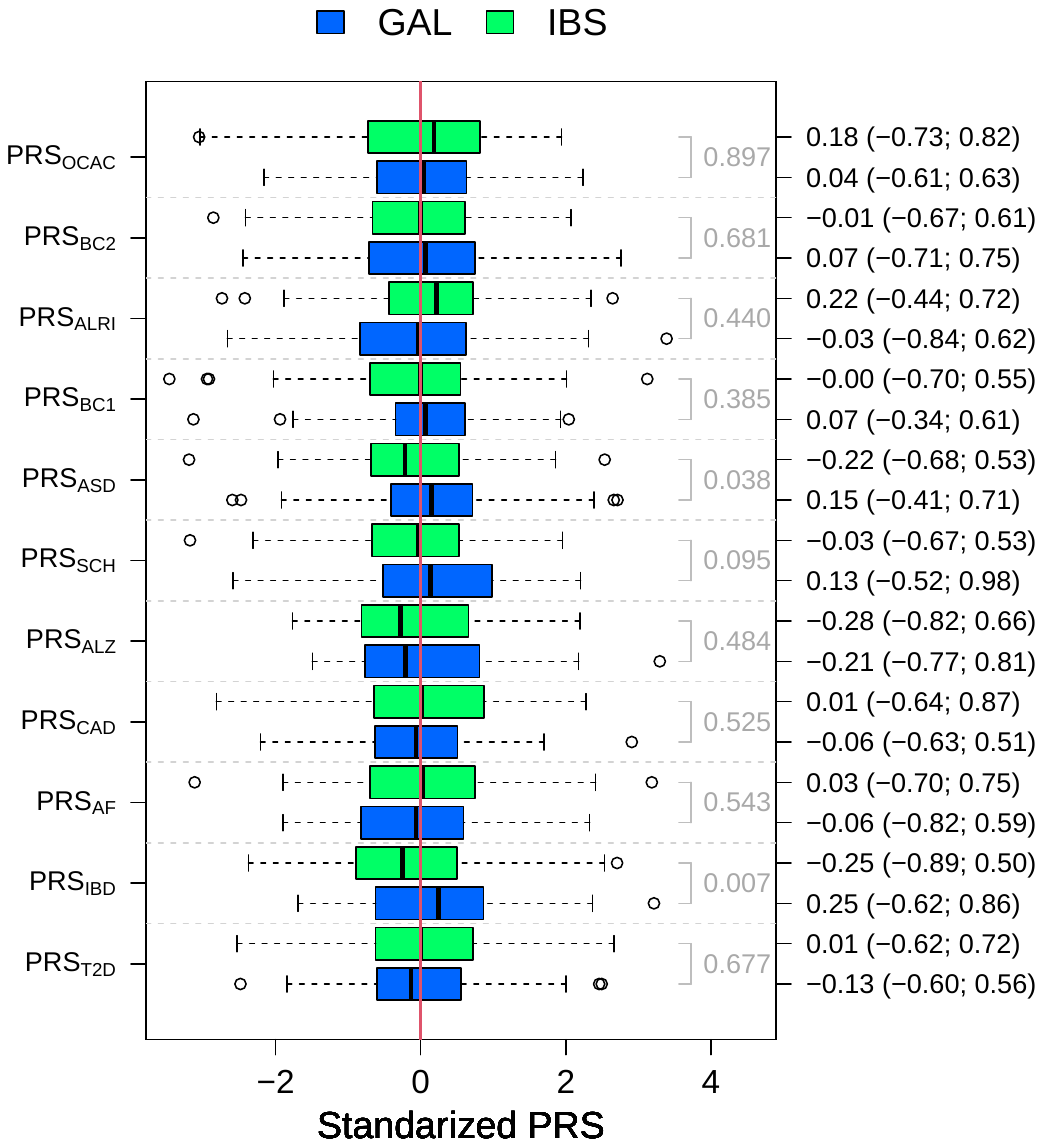

**Figure S7** Boxplots of PRS values for different common diseases within Galician fineSTRUCTURE clusters.

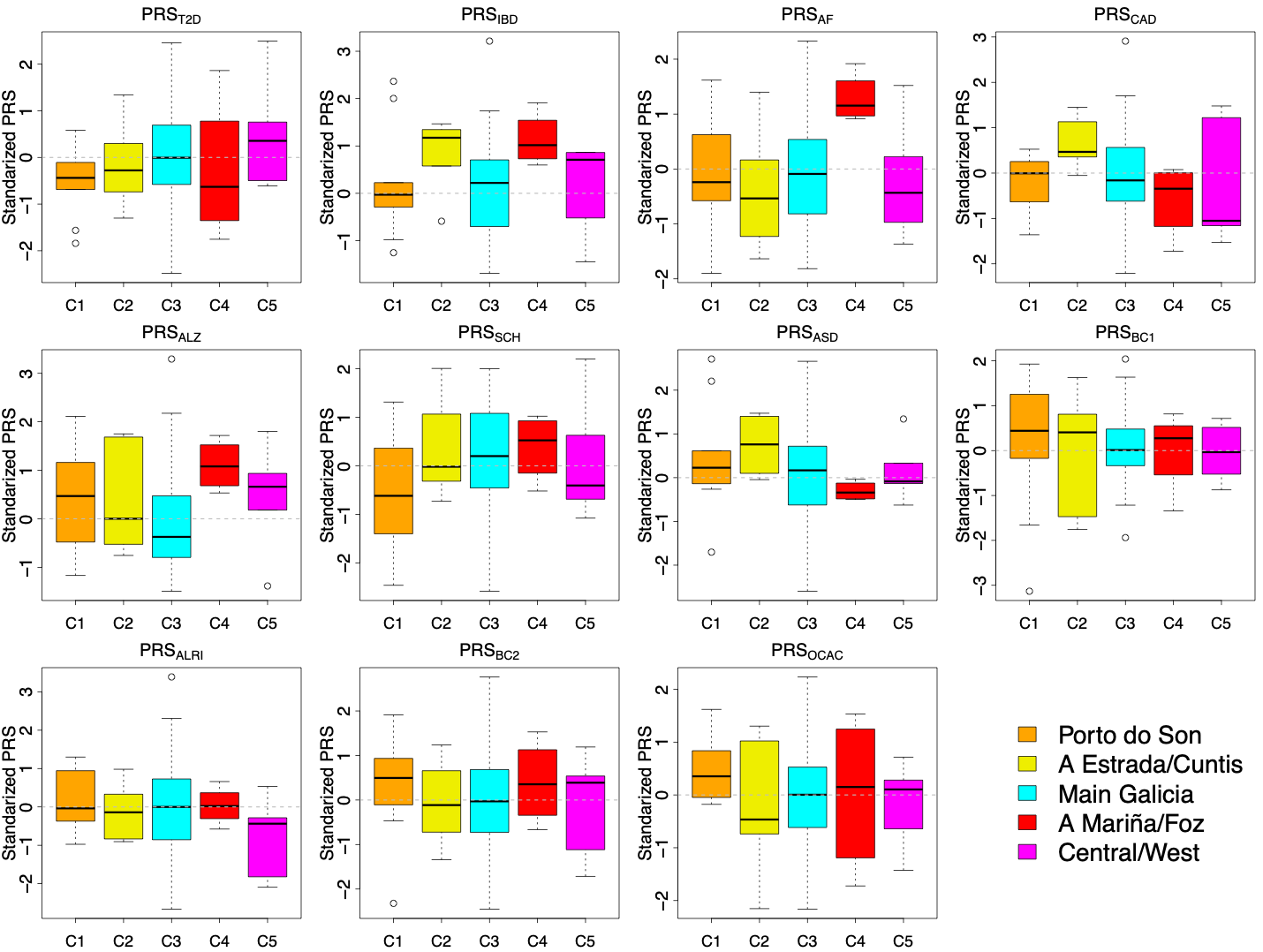
