## Supplementary Text for "Whole-genome sequencing in Galicia reveals male-biased pre-Islamic North African ancestry, subtle population structure, and micro-geographic patterns of disease risk"

**Supplementary Text 2**

**About North African ancestry in Iberia**

Several studies have consistently reported a significant North African genetic component among Iberians, reflecting the historical influence of the Moorish invasion of the Peninsula in the 8^th^ century. Many of these findings are based on the analysis of uniparental markers (Adams *et al.*, 2008; Capelli *et al.*, 2009; Moorjani *et al.*, 2011; Semino *et al.*, 2004). For mitochondrial DNA (mtDNA), the North African contribution, primarily measured by the frequency of the U6 haplogroup, ranges between 0% to 3.7% (Barral-Arca *et al.*, 2016) in different continental Iberian regions (e.g. Andalusia: 2.9% [with a peak in the province of Huelva: ~12%]; Galicia: 1.1%; Murcia: 3.7%; inferred from their Table S1 and Table S2 (Barral-Arca *et al.*, 2016)), with other report reaching up to 8.5% (Pereira *et al.*, 2005). For the Y-chromosome, estimates vary significantly across regions, reported values include ranges of: 2.1–9.2% (Andalucia: 9.2%; Basques: 2.1%; Catalonia: 6.1%) (Bosch *et al.*, 2001), 2.3–20.8% (Adams *et al.*, 2008) (Andalucia East: 2.4%; Andalucia West, 16.7%; Asturias: 10.5%; Castile Northwest: 21.7%; Castile Northeast, 9.3%; Catalonia: 2.3%; Extremadura: 19%; Galicia, 20.8%; Valencia: 12.8%), 1.5–18% (Capelli *et al.*, 2009) (Andalucia: 5.4%; Basques: 1.5%; Cantabria: 18%; Galicia: 6.8%; Spain: 7.7%) to 2.1–9.2%(Semino *et al.*, 2004) (Andalucia: 9.2%; Basques: 2.1%; Catalonia: 6.1%). Estimates based on autosomal DNA provide further insight into the North African component in Iberia. For continental Spain, values range from 7–9% (Botigué *et al.*, 2013) (Andalucia: 9%; Basques: 7%; Galicia: 9%; Spain Central: 9%). Other autosomal analyses report values ranging from 0–11% (Bycroft *et al.*, 2019)(Basques: ~0%; West Peninsula [excluding Galicia]: ~10%; Galicia: 11%). The presence of a North African component in Iberia has also been documented in ancient DNA sequencing studies. Silva et al. (2021) identified evidence of admixture between Amazigh (Berber) people and the local population inhabiting the Iberian Peninsula prior to the Islamic conquest, based on the analysis of an eleventh-century CE man buried in an Islamic necropolis in Segorbe (Valencia, located at the Southeast Mediterranean coast in Spain). Similarly, Olalde et al. (2019) revealed intermittent pre-Roman interactions between North Africa and Iberia, as well as substantial gene flow of North African ancestry into Iberia arising around the Roman period.

**References**

Adams, S. M., Bosch, E., Balaresque, P. L., Ballereau, S. J., Lee, A. C., Arroyo, E.*, et al.* (2008). The genetic legacy of religious diversity and intolerance: paternal lineages of Christians, Jews, and Muslims in the Iberian Peninsula. *Am J Hum Genet* **83,** 725-36.

Barral-Arca, R., Pischedda, S., Gómez-Carballa, A., Pastoriza, A., Mosquera-Miguel, A., López-Soto, M.*, et al.* (2016). Meta-Analysis of Mitochondrial DNA Variation in the Iberian Peninsula. *PLoS One* **11,** e0159735.

Bosch, E., Calafell, F., Comas, D., Oefner, P. J., Underhill, P. A. &Bertranpetit, J. (2001). High-resolution analysis of human Y-chromosome variation shows a sharp discontinuity and limited gene flow between northwestern Africa and the Iberian peninsula. *Am. J. Hum. Genet.* **68,** 1019-1029.

Botigué, L. R., Henn, B. M., Gravel, S., Maples, B. K., Gignoux, C. R., Corona, E.*, et al.* (2013). Gene flow from North Africa contributes to differential human genetic diversity in southern Europe. *Proc Natl Acad Sci U S A* **110,** 11791-6.

Bycroft, C., Fernandez-Rozadilla, C., Ruiz-Ponte, C., Quintela, I., Carracedo, A., Donnelly, P.*, et al.* (2019). Patterns of genetic differentiation and the footprints of historical migrations in the Iberian Peninsula. *Nat Commun* **10,** 551.

Capelli, C., Onofri, V., Brisighelli, F., Boschi, I., Scarnicci, F., Masullo, M.*, et al.* (2009). Moors and Saracens in Europe: Estimating the medieval North African male legacy in southern Europe. *Eur. J. Hum. Genet.* **17,** 848-52.

Moorjani, P., Patterson, N., Hirschhorn, J. N., Keinan, A., Hao, L., Atzmon, G.*, et al.* (2011). The history of African gene flow into Southern Europeans, Levantines, and Jews. *PLoS Genet* **7,** e1001373.

Olalde, I. & Mallick, S. & Patterson, N. & Rohland, N. & Villalba-Mouco, V. & Silva, M.*, et al.* (2019). The genomic history of the Iberian Peninsula over the past 8000 years. *Science* **363,** 1230-1234.

Pereira, L., Cunha, C., Alves, C. &Amorim, A. (2005). African female heritage in Iberia: a reassessment of mtDNA lineage distribution in present times. *Hum Biol* **77,** 213-29.

Semino, O., Magri, C., Benuzzi, G., Lin, A. A., Al-Zahery, N., Battaglia, V.*, et al.* (2004). Origin, diffusion, and differentiation of Y-chromosome haplogroups E and J: Inferences on the neolithization of Europe and later migratory events in the Mediterranean area. *Am. J. Hum. Genet.* **74,** 1023-34.

Silva, M., Oteo-Garcia, G., Martiniano, R., Guimaraes, J., von Tersch, M., Madour, A.*, et al.* (2021). Biomolecular insights into North African-related ancestry, mobility and diet in eleventh-century Al-Andalus. *Sci Rep* **11,** 18121.
